# Establishing Conserved Biosynthetic Gene Clusters of the Phylum Myxococcota

**DOI:** 10.1101/2025.06.19.660557

**Authors:** Shailaja Khanal Pokharel, Nawal Shehata, Andrew Ahearne, Thomas Knehans, Constance B. Bailey, Paul D. Boudreau, D. Cole Stevens

## Abstract

A surge in sequenced myxobacteria catalyzed by advancements in long read genome and metagenome sequencing has provided sufficient data to scrutinize the conserved biosynthetic gene clusters (BGCs) within the phylum Myxococcota. Provided the utility of myxobacteria in environmental nutrient cycles and discovery of novel therapeutic leads, we sought to determine any conserved specialized metabolism in the phylum. Using a pan-genome approach to analyze eleven genera and 195 sequenced genomes including ten newly reported myxobacterial isolate, we observed five conserved BGCs. All five clusters encode for characterized metabolites with established ecological roles for four of the metabolites, and none of the metabolites are known toxins. Validation of our approach was done by analyzing Myxococcota genera without sufficient, sequenced representatives for pan-genome analysis to observe the presence/absence of these five clusters. This approach enabled observation of genus-level conservation of BGCs with varying degrees of confidence due to diversity of sequenced species within each genus. The indigoidine BGC typically found in *Streptomyces* spp. was notably conserved in *Melittangium*; heterologous expression of the core biosynthetic gene *bspA* in *Escherichia coli* and subsequent detection of indigoidine confirmed the identity of the indigoidine cluster. Conserved BGCs in myxobacteria reveal maintenance of biosynthetic pathways and cognate metabolites with ecological roles as chemical signals and stress response; these observations suggest competitive specialization of secondary metabolism and toxin production in myxobacteria.

## Intro

Myxococcota possess numerous features that motivate continued discovery of environmental isolates such as cosmopolitan, predatory lifestyles that contribute to soil nutrient cycles and extraordinary capacities to produce specialized metabolites with potential applications as therapeutic leads (1-5). A recent expansion of the phylum brought about by isolation and characterization of novel species and genera has generated a wealth of genomic data to be scrutinized. Bioinformatic analyses has previously revealed all genera within the phylum, excluding *Vulgatibacter* (6), accommodate numerous biosynthetic gene clusters (BGCs) responsible for the production of specialized metabolites (7, 8), and metabolomic analysis has demonstrated a correlation between metabolic profiles and taxonomic distance of myxobacteria (9). Common BGCs repeatedly observed in myxobacterial genomes have been noted such as the geosmin BGC (10). However, the distribution and conservation of specific BGCs from myxobacteria have not been reported. Establishing a conserved set of BGCs across the Myxococcota will improve dereplication of common clusters for future genome mining and inform efforts to determine the ecological role of specialized metabolites from myxobacteria.

Inspired by a pan-genome analysis of *Corallococcus* spp. that found the core pan-genome excluded the majority of BGCs observed in the genus (11), we sought to determine the distribution and conservation of BGCs from sequenced Myxococcota. Although numerous database-driven pipelines for BGC analysis exist to annotate biosynthetic genes and determine pathway novelty (12, 13), we opted for an initial pan-genome analysis to identify conserved genes that were subsequently mapped back to BGCs annotated by antiSMASH (14, 15). Using an approach that is initially BGC-agnostic, we hoped to minimize challenges associated with ambiguous similarity scores of BGCs across analysis pipelines, influence of intrinsic homologies of core biosynthetic features, and occurrences of multiple discrete clusters in individual BGCs annotated as “hybrids.” We analyzed a total of 195 myxobacterial genomes from eleven genera and found five BGCs to be generally conserved in myxobacteria.

## Results

### Genome sequencing and phylogenetic analysis of 10 isolated myxobacteria

Myxobacterial isolates were obtained from rhizospheric soil samples using prey-baiting methodology as previously described (16, 17). A total of ten environmental isolates were sequenced to broaden our pan-genome analysis (Table 1). Taxonomic assignments for sequenced isolates included three *Archangium*, one *Cystobacter*, two *Melittangium*, two *Myxococcus*, and two *Nannocystis*. Comparative genome analysis of newly obtained isolates versus type strain myxobacteria provided average nucleotide identity (ANI) and digital DNA– DNA hybridization values (dDDH) supported several isolates being novel species (Table 1). Adhering to established thresholds for determining novel species (18, 19), isolates PVMSAZ, D1P2, TKBC04, and BB12-2 are candidate novel species. However TKBC04 and D1P2 are both the same species and share 99.9% ANI. Both isolates also share >97% ANI with *Melittangium primigenium*. We previously reported the misassignment of *Me. primigenium* as *Archangium primigenium* ATCC 29037 and feel compelled to note the candidate status of the novel species represented by strains D1P2 and TKBC04 until further effort is done to validly describe and resolve the type strain for these *Melittangium* (17). The low-quality assembly of BB12-2 (>100 contigs) decreases our confidence in it being a novel *Myxococcus* species. Of the three *Archangium* isolates, PVMSAZ, a candidate novel species, is most related to *Archangium gephyra* (90.6% ANI), NCHinoki1 is a subspecies of *Archangium lansingense*, and SCPoplar1 is a subspecies of *Ar. gephyra*. Isolates NCWS, and Hickory4 are subspecies of *Corallococcus terminator, Cystobacter fuscus*, and *Myxococcus fulvus* respectively. Isolates MIELM and UBH4 are both subspecies of *Nannocystis pusilla*.

**Table 1.** Genome assembly and phylogenetic data for sequenced isolates with proposed novel species bolded. ^*^Isolates *Me*. D1P2 and *Me*. TKBC04 are the same species and share 99.9% ANI. *Myxococcus llanfairpwllgwyngyllgogerychwyrndrobwllllantysiliogogogochensis* abbreviated to *Myxococcus llanfair*….

| Isolate | Size (bp) | CDS | GC% | N50 | L50 | contigs | dDDH (d4) | ANI |
| --- | --- | --- | --- | --- | --- | --- | --- | --- |
| *Archangium* sp. NCHinoki1 | 13,608,435 | 11,298 | 68.2 | 13,506,606 | 1 | 2 | 35.5%; *Archangium gephyra* | 98.7%; *Archangium lansingense* |
| ***Archangium* sp. PVMSAZ** | **10,952,989** | **9,658** | **69.6** | **10,843,764** | **1** | **2** | **40.6%; *Ar. gephyra*** | **90.6%; *Ar. gephyra*** |
| *Archangium* sp. SCPoplar1 | 13,446,571 | 10,895 | 68.7 | 13,291,844 | 1 | 2 | 45.6%; *Ar. gephyra* | 95.8%; *Ar. gephyra* |
| *Cystobacter* sp. NCWS | 12,322,754 | 14,750 | 68.6 | 12,003,056 | 1 | 3 | 60.8%; *Cystobacter fuscus* | 95.2%; *Cy. fuscus* |
| ***Melittangium* sp. D1P2\*** | **9,298,785** | **7,782** | **70.8** | - | **1** | **1** | **29.5%; *Melittangium boletus*** | **85.7%; *Me. boletus*** |
| ***Melittangium* sp. TKBC04\*** | **9,298,954** | **7,763** | **70.8** | - | **1** | **1** | **29.5%; *Me. boletus*** | **85.7%; *Me. boletus*** |
| ***Myxococcus* sp. BB12-2** | **10,892,721** | **9,003** | **69** | **130,546** | **23** | **148** | **30.2%; *Myxococcus llanfair...*** | **85.2%; *My. llanfair...*** |
| *Myxococcus* sp. Hickory4 | 10,160,340 | 8,489 | 70.1 | 108,358 | 30 | 182 | 59.7%; *Myxococcus fulvus* | 95.1%; *My. fulvus* |
| *Nannocystis* sp. MIELM | 12,731,429 | 14,925 | 71.4 | 12,694,743 | 1 | 3 | 56.6%; *Nannocystis pusilla* | 94.9%; *Na. pusilla* |
| *Nannocystis* sp. UBH4 | 12,213,092 | 11,265 | 71.7 | - | 1 | 1 | 55.4%; *Na. pusilla* | 94.7%; *Na. pusilla* |

### Pan-genome analysis of Myxococcota

Pan-genomes for each Myxococcota genus were determined from a total of 195 genome assemblies with available genomes per genus ranging from 63 *Myxococcus* genomes to four *Melittangium* genomes (Figure 1). Genera excluded from our analysis due to an insufficient number of sequenced representatives include *Aggregicoccus, Chondromyces, Citreicoccus, Enhygromyxa, Haliangium, Hyalangium, Kofleria, Labilithrix, Plesiocystis, Pseudenhygromyxa, Sandaracinus, Simulacricoccus, Vitiosangium*, and *Vulgatibacter*. Pan-genomes for members of the following genera were determined *Myxococcus, Corallococcus, Anaeromyxobacter, Sorangium, Archangium, Nannocystis, Polyangium, Pyxidicoccus, Cystobacter, Stigmatella*, and *Melittangium* (Figure 1). Our analysis suggests all analyzed genera have open genomes with the majority of genes found to be in the accessory pan-genomes (shell and cloud). Clades with higher percentages of genes in core pan-genomes (*Melittangium* 6.8% and *Stigmatella* 14%) have fewer sequenced genomes available, and ANI values for members indicate these genera include genome data from highly related sub-species and fewer distinct species. For example, of the four *Melittangium* analyzed, only two distinct species with ANI and dDDH values below established thresholds are included in our analysis. The limited diversity of species in the *Cystobacter, Melittangium*, and *Stigmatella* clades likely impacts our assessment of conserved BGCs at the genus-level. Despite this limitation, we retained and included all three genera with the caveat that the following BGCs conserved at the genus-level for the *Cystobacter, Melittangium*, and *Stigmatella* may be overstated.

**Figure 1.**
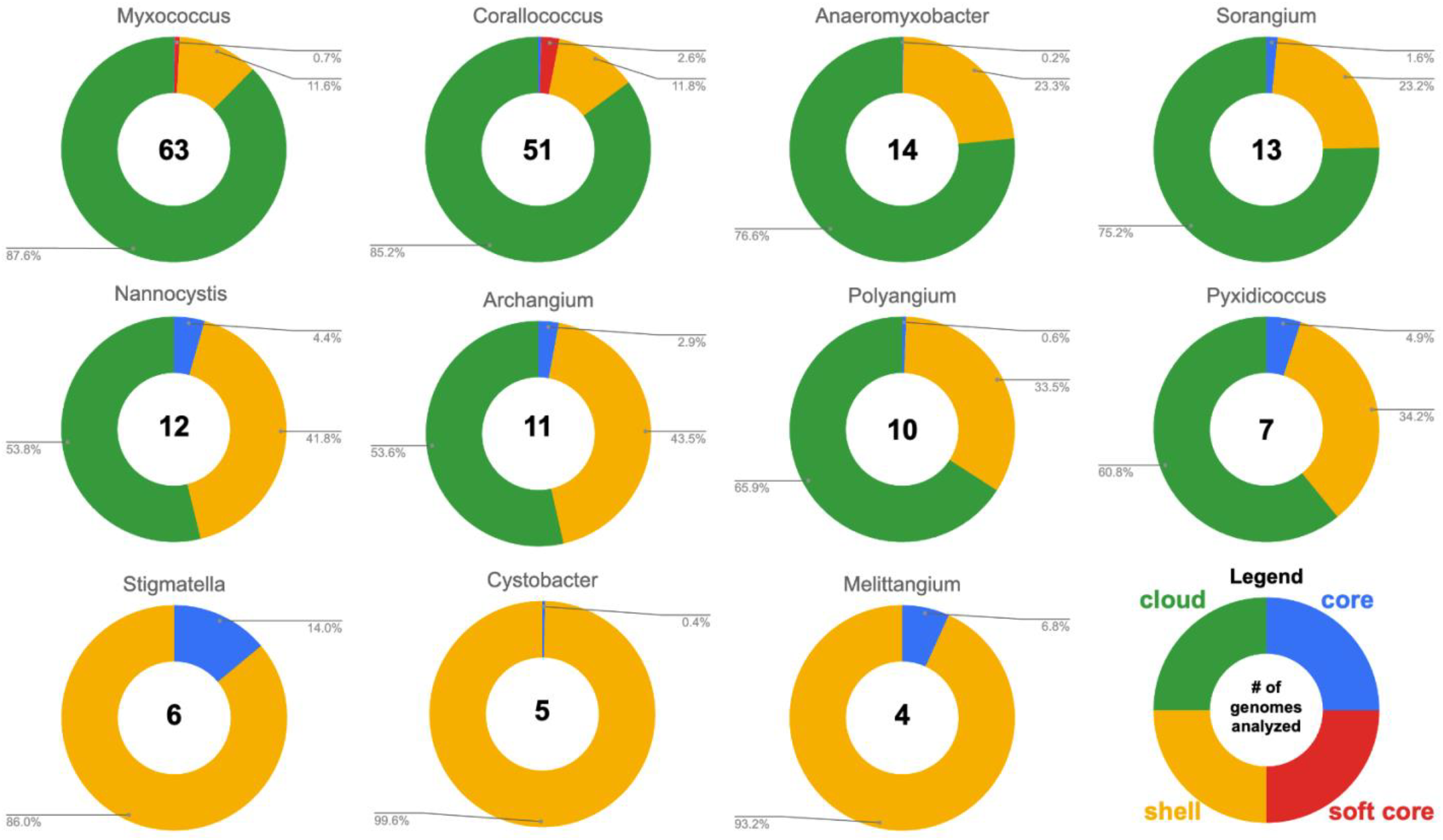
Summary of pan-genome analysis by genera. A complete list of myxobacteria included in our analysis can be found as supplemental data (Table S1).

### Identification of five conserved BGCs in myxobacteria

Pan-genome results were used to determine genes observed in ≥50% of the total number of analyzed genomes per genus. The identified genes were then mapped to BGCs predicted from each assembly using antiSMASH. This process provided lists of genes conserved in ≥50% of analyzed genomes per genus and their presence/absence in BGCs from each genome. Clusters present in ≥50% of the total number of analyzed genomes per genus that included ≥2 genes from resulting lists were considered conserved at the genus-level. Conserved BGCs from each genus were aggregated and used to assess the distribution of common BGCs within the phylum. AntiSMASH similarity scores and comparison with representative BGCs in the MiBIG database afforded assignments for characterized BGCs and associated metabolites (20, 21).

Five BGCs are present in at least seven of the eleven genera analyzed (Table 2). All but one of these five BGCs can be unambiguously assigned using validated BGCs deposited in the MiBIG database including the geosmin (BGC0001181), carotenoid (BGC0000648), VEPE/AEPE/TG-1 (BGC0000871), and myxochelin (BGC0002492) clusters (22-29). The remaining cluster has varying similarity (20-63%) with a characterized alkylpyrone BGC (BGC0001831) and includes core type III polyketide synthase (PKS) features for alkylpyrone production (30). Maintenance of cluster organization and ANIs for each BGC was obtained using clinker and OAT respectively to assess the validity of our approach (Figures 2-4) (31-33). This revealed shared identities for biosynthetic gene products and co-located gene products not anticipated to be directly involved in metabolite assembly. For example, recently discovered encapsulins thought to be involved in small, volatile terpene trafficking are consistently present in all geosmin clusters (34, 35). Maintained spatial organization of the five clusters can be observed at the genus-level, and organization of the VEPE/AEPE/TG-1 and carotenoid clusters is especially conserved in Myxococcia (Figures 2 and 3). The VEPE/AEPE/TG-1 cluster is the only characterized cluster conserved in *Anaeromyxobacter*, and the geosmin cluster is the only characterized cluster conserved in *Nannocystis*. The VEPE/AEPE/TG-1 and alkylpyrone clusters are not conserved in any analyzed Polyangiia (*Nannocystis, Polyangium*, and *Sorangium*). Interestingly, our analysis suggests a gene duplication event apparent in the alkylpyrone BGCs from *Archangium* may have contributed to the differences in cluster organization in the Myxococcia (Figure 3). The consistent presence, organization, and high identities between these five clusters are indicative of common ancestry and vertical inheritance in the phylum.

**Table 2.** Genus-level presence/absence of the five BGCs observed to be conserved in Myxococcota.

| BGC | genera with | genera without |
| --- | --- | --- |
| geosmin | *Archangium*, *Corallococcus*, *Cystobacter*, *Melittangium*, *Myxococcus*, *Nannocystis*, *Polyangium*, *Pyxidicoccus*, *Sorangium*, *Stigmatella* | *Anaeromyxobacter* |
| carotenoid | *Archangium*, *Corallococcus*, *Cystobacter*, *Melittanigum*, *Myxococcus*, *Polyangium*, *Pyxidicoccus*, *Sorangium*, *Stigmatella* | *Anaeromyxobacter*,<br>*Nannocystis* |
| VEPE/AEPE/TG-1 | *Anaeromyxobacter*, *Archangium*, *Corallococcus*, *Cystobacter*, *Melittanigum*, *Myxococcus*, *Pyxidicoccus*, *Stigmatella* | *Nannocystis*,<br>*Polyangium*, *Sorangium* |
| alkylpyrone | *Archangium*, *Corallococcus*, *Cystobacter*, *Melittanigum*, *Myxococcus*, *Pyxidicoccus*, *Stigmatella* | *Anaeromyxobacter*,<br>*Nannocystis*,<br>*Polyangium*, *Sorangium* |
| myxochelin | *Archangium*, *Corallococcus*, *Cystobacter*, *Melittanigum*, *Myxococcus*, *Pyxidicoccus*, *Polyangium*, *Stigmatella* | *Anaeromyxobacter*,<br>*Nannocystis*, *Sorangium* |

**Figure 2.**
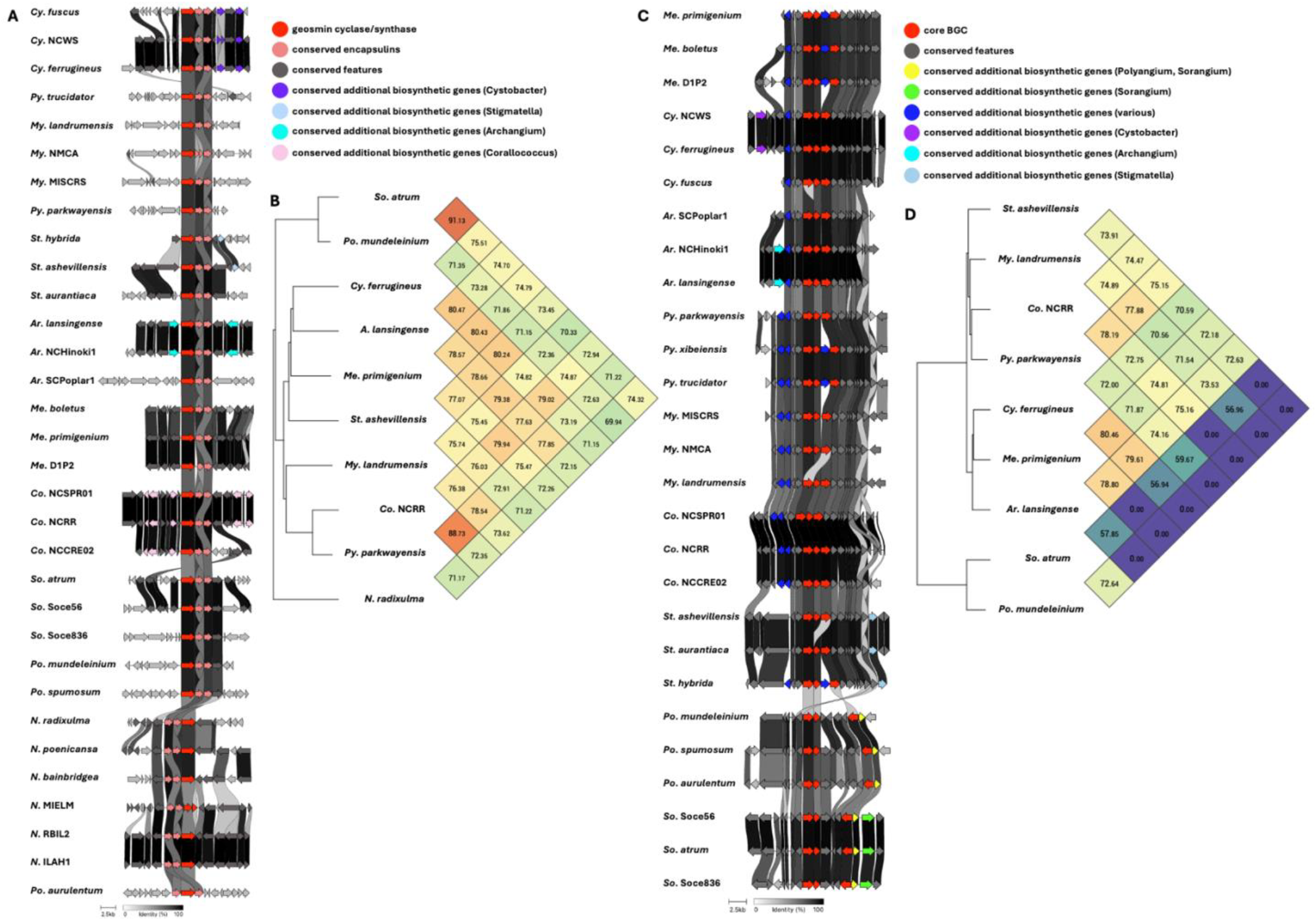
Conservation and genetic organization of the geosmin BGC in Myxococcota (A) and ANI values for geosmin BGCs across representatives of host genera (B). Conservation and genetic organization of the carotenoid BGC in Myxococcota (C) and ANI values for carotenoid BGCs across representatives of host genera (D). AntiSMASH analysis provided .gbk files utilized to generate images in clinker and ANI values using OrthoANI (32, 64).

**Figure 3.**
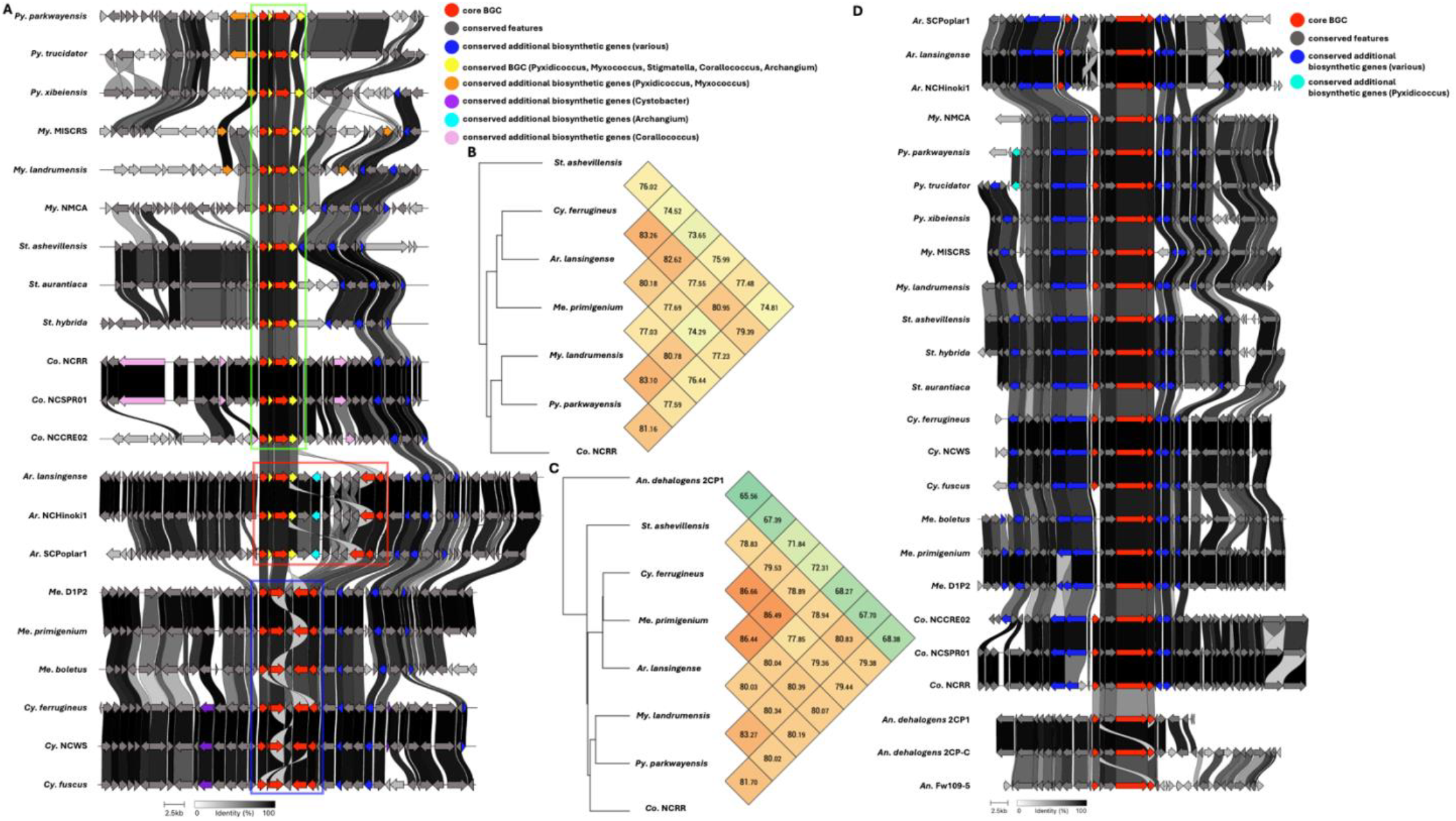
Conservation and genetic organization of the alkylpyrone BGC in Myxococcota (A) and ANI values for alkylpyrone BGCs across representatives of host genera (B). The discussed gene duplication event in *Archangium* (red box) and resulting differences in *Corallococcus, Pyxidicoccus, Myxococcus* (green box) and *Cystobacter* and *Melittangium* (blue box) clusters are indicated. Conservation and genetic organization of the VEPE/AEPE/TG-1 BGC in Myxococcota (D) and ANI values for VEPE/AEPE/TG-1 BGCs across representatives of host genera (C). AntiSMASH analysis provided .gbk files utilized to generate images in clinker and ANI values using OrthoANI (32, 64).

### Validation of identified BGCs conserved in myxobacteria

We sought to test the validity of our observation that the geosmin, carotenoid, VEPE/AEPE/TG-1, alkylpyrone, and myxochelin clusters are conserved in myxobacteria by determining presence/absence of each BGC from lesser-studied genera not included in our pan-genome analysis due to lack of sequenced representatives (Table 3). Notably, *Kofleria* and *Simulacricoccus* had no sequenced species to include in our analysis, and poor genome quality precluded the only sequenced *Plesiocystis, Plesiocystis pacifica* (36). At least two of five conserved BGCs were present in all myxobacteria included. All five BGCs were present in representative *Citreicoccus, Hyalangium*, and *Vitiosangium*. Representatives from *Labilithrix, Pseudenhygromyxa*, and *Vulgatibacter* host the fewest conserved clusters with only two of the five BGCs present in each. However, it is worth noting *Vulgatibacter incompetus* has just four BGCs in total, so the conserved VEPE/AEPE/TG-1 and alkylpyrone clusters present account for half of the BGCs in the genome. The myxochelin BGC was the least conserved cluster observed in the eleven analyzed myxobacteria (5/11), and the carotenoid and alkylpyrone BGCs were the most conserved (9/11).

**Table 3.**
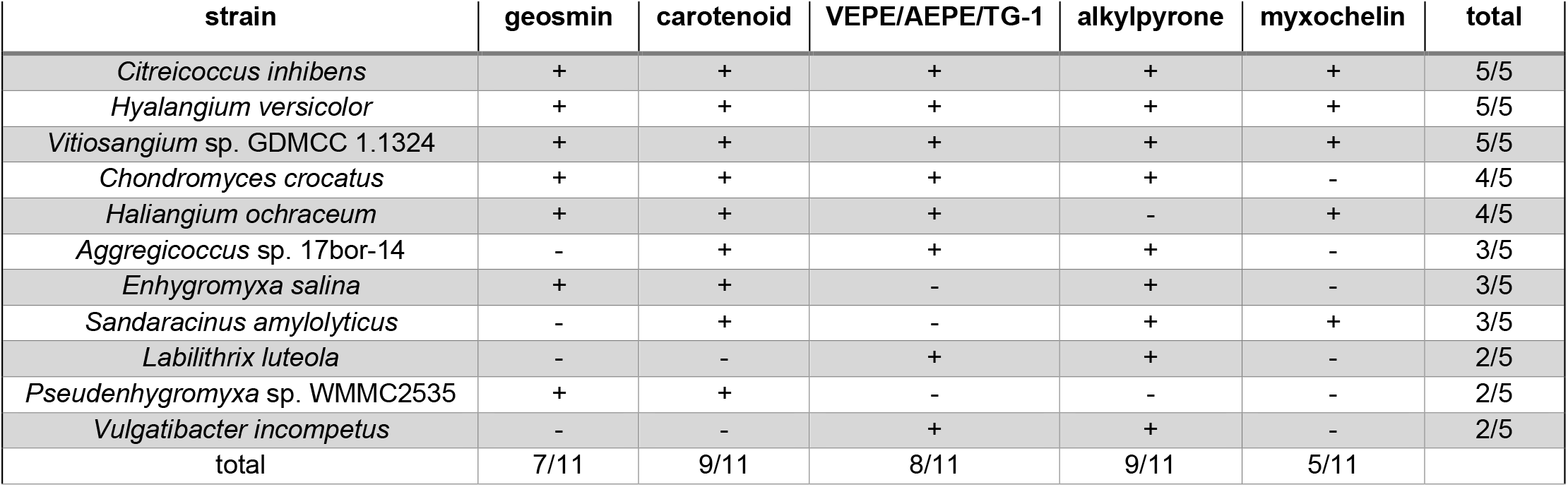
Validation of conserved BGCs using presence/absence data of representatives from genera not included in our pan-genome analysis.

### Genus-level conservation of BGCs in myxobacteria

Using our genus-level pan-genome data, we were able to catalog and compare the distribution of BGCs across genera included in our dataset (Table 4). The five BGCs conserved at the phylum-level are the only characterized BGCs conserved in *Corallococcus* and *Cystobacter*. Beyond the five assigned BGCs conserved in the phylum, only two characterized BGCs are conserved across multiple genera. The myxoprincomide BGC (BGC0000393) is conserved in *Myxococcus* and *Pyxidicoccus* (37), and the dkxanthene BGC (BGC0000986) is conserved in *Myxococcus* and *Stigmatella* (38). The indigoidine BGC (BGC0000727) from *Streptomyces aureofaciens* is conserved in *Melittangium* (39). The indigoidine BGC is also present in *Cystobacter ferrugineus* and *Polyangium* sp. y55×31. Shared identities with indigoidine clusters from *Str. aureofaciens* and *Streptomyces chromofuscus* (BGC0000375) suggest horizontal transfer of the cluster (Figure 5A) (40). Excluding *Polyangium*, uncharacterized BGCs that encode for unknown metabolites are conserved in all genera (Table 4). Of these, a type 1 PKS cluster conserved in *Archangium* and *Melittangium* was the only unknown BGC present in multiple genera.

**Table 4.**
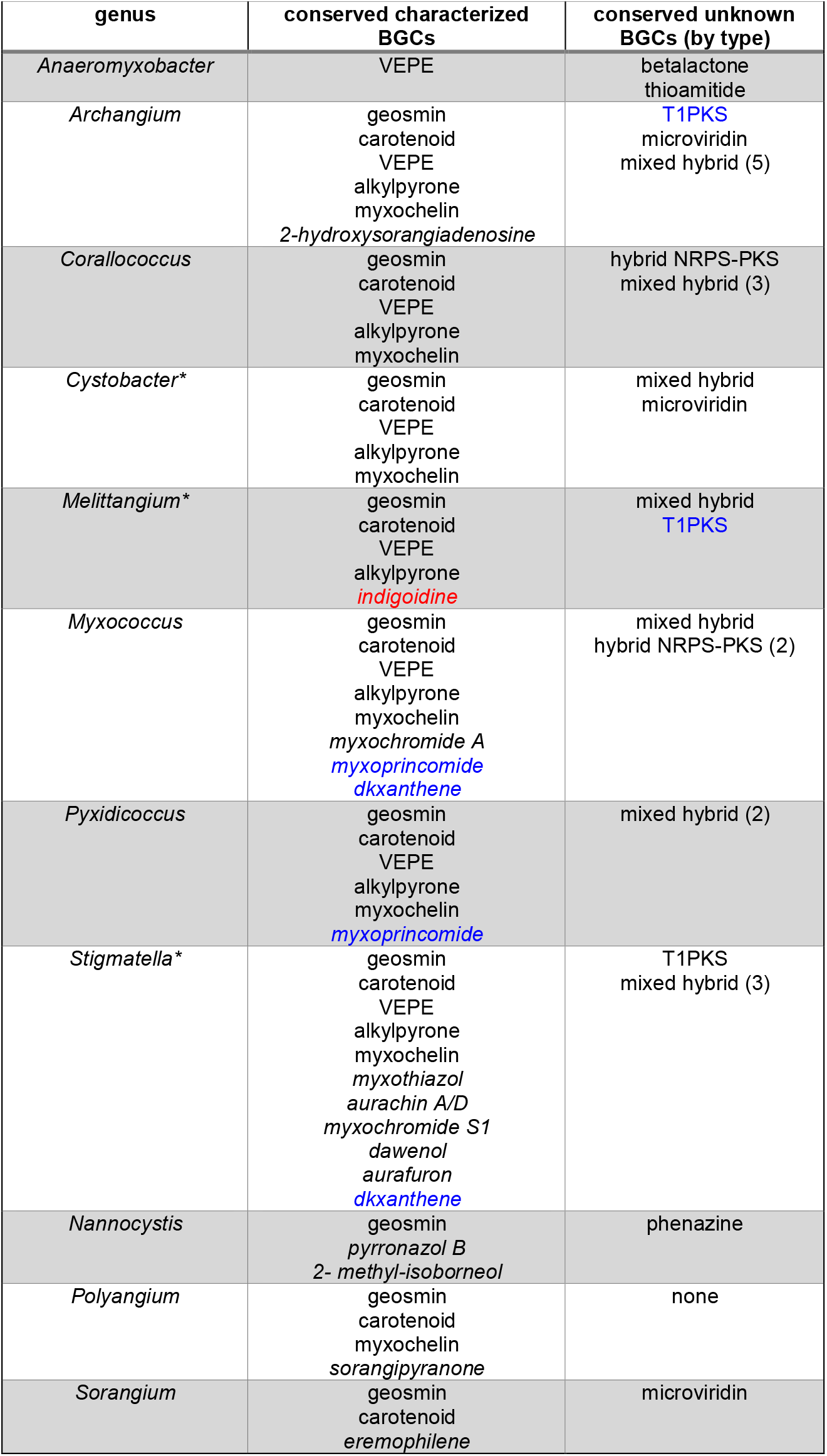
Conserved characterized (top) and uncharacterized (bottom) BGCs identified by pan-genome analysis organized by genus. Clusters conserved at the genus-level that are not broadly conserved in the phylum are italicized. AntiSMASH-assigned cluster types were used to label uncharacterized BGCs with totals for each type included in parentheses when necessary. BGCs present in multiple genera are colored blue. *Genera with fewer sequenced distinct species and lower confidence of cluster conservation.

**Figure 4.**
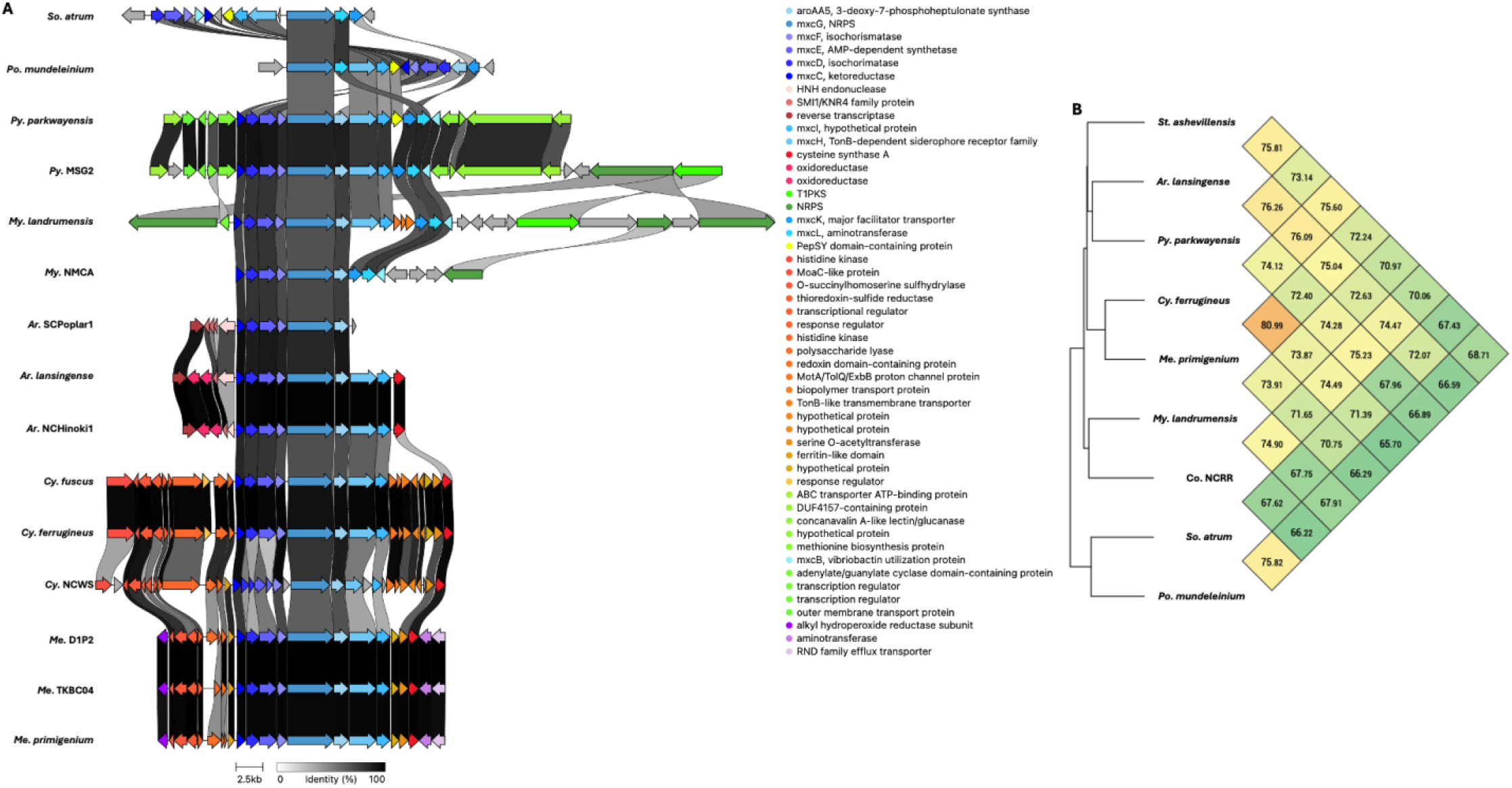
Conservation and genetic organization of the myxochelin BGC in Myxococcota (A) and ANI values for myxochelin BGCs across representatives of host genera (B). AntiSMASH analysis provided .gbk files utilized to generate images in clinker and ANI values using OrthoANI (32, 64).

**Figure 5.**
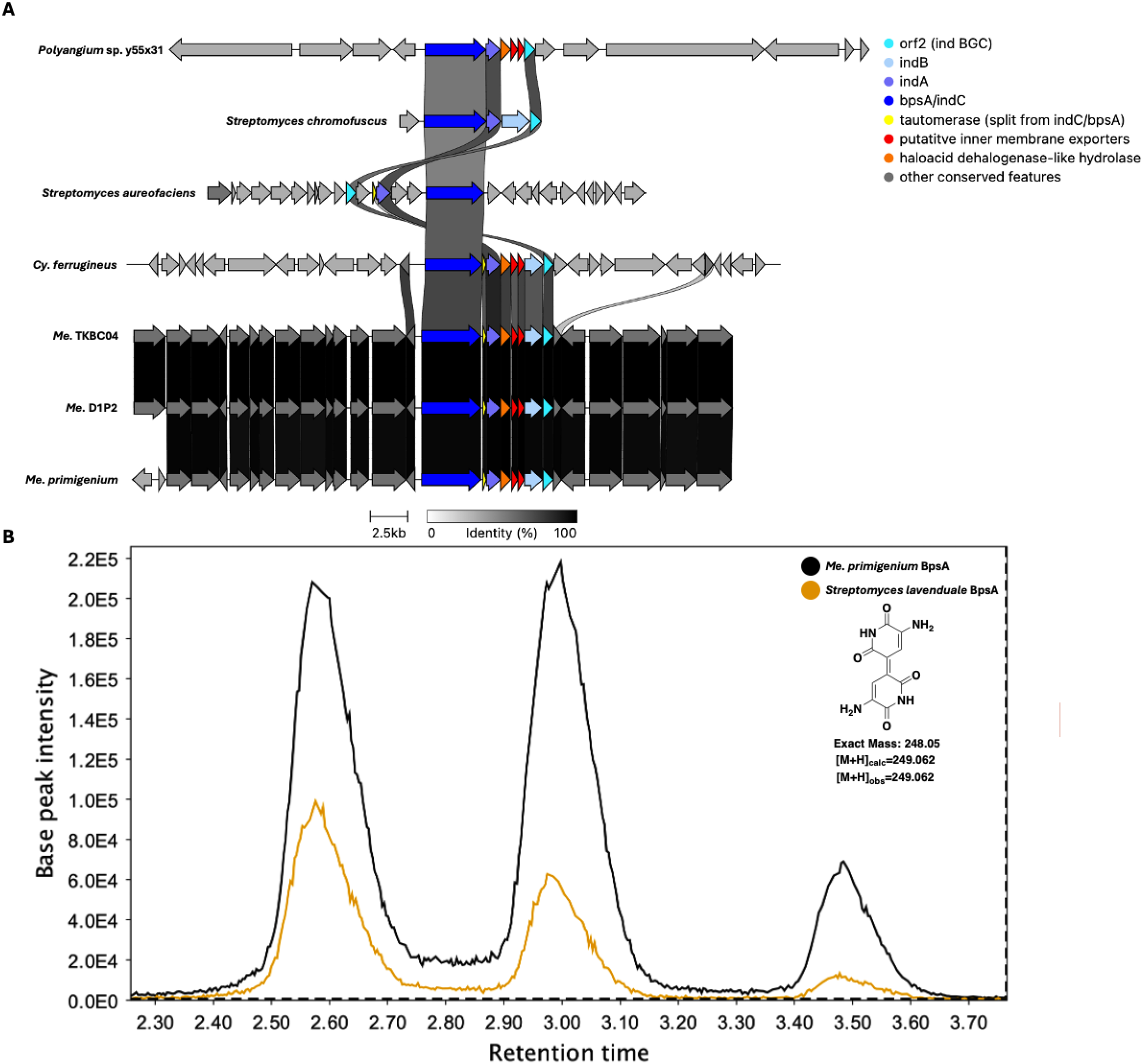
Conservation and genetic organization of the indigoidine BGC from *Str. chromofuscus, Str. aureofaciens*, and host Myxococcota (A). Extracted ion chromatograms (EICs) for 249-249.1 m/z obtained by LC-MS/MS analysis of extracts from *E. coli* BAP1 expressing *bpsA* homologs from *Str. lavenduale* and *Me. primigenium*. AntiSMASH analysis provided .gbk files for indigoidine clusters of Myxococcota (64). Indigoidine clusters for *Streptomyces* species were obtained from the MIBIG database as .gbk files. MZMine was used to generate EIC images (20).

### Heterologous expression of BpsA from *Me. primigenium*

An *indC*/*bpsA* homolog (WP_204491470.1) encoding the nonribosomal synthetase (NRPS), blue pigment synthetase A (BpsA), in the identified indigoidine BGC from *Me. primigenium* was heterologously expressed in *Escherichia coli* BAP1 to confirm similarity-based assignment of the cluster. Numerous heterologous platforms for indigoidine production have been developed by expression of BpsA in various hosts (41-44). Comparing *E. coli* BAP1 expressing BpsA from *Me. primigenium* with *E. coli* BAP1 expressing BpsA from *Streptomyces lavendulae* (45, 46), an established indigoidine producing strain (47, 48), we observed blue phenotypes indicative of indigoidine production from each strain. Successful indigoidine production of *E. coli* BAP1 expressing *indC*/*bpsA* from *Me. primigenium* was confirmed by subsequent LC-MS/MS analysis of organic phase extracts from both *E. coli* strains comparing indigoidine retention times, *m/z*, and fragmentation pattern (Figure 5B). These results confirm the similarity-based assignment and noted conservation of the indigoidine BGC in *Melittangium*.

## Discussion

We identified five clusters using our pan-genome approach to identify BGCs conserved in myxobacteria. The high level of conservation and spatial organization found in each of the five conserved BGCs suggests vertical inheritance from a common ancestor. Vertical inheritance of these five clusters is also supported by variable ANIs for clusters present in extant lineages. Corroborating an inspirational pan-genome analysis of *Corallococcus* (11), we also found the majority of myxobacterial BGCs to be excluded from core pan-genomes for all analyzed genera. As noted, the low availability of representative, distinct species of *Cystobacter, Melittangium*, and *Stigmatella* offers a clear limitation in our approach. Validation of conservation for these five BGCs by determining their presence in available genome data of representative myxobacteria from genera not included in our pan-genome analysis revealed variable conservation for each cluster. These data support our conclusion that the geosmin, carotenoid, VEPE/AEPE/TG-1, myxochelin, and alkylpyrone clusters are broadly conserved in Myxococcota.

Excluding alkylpyrones, metabolites produced from all other conserved BGCs have characterized ecological functions. Geosmin is a ubiquitous, volatile sesquiterpenoid produced by numerous prokaryotes and eukaryotes (34). Geosmin producers repel predatory *Caenorhabditis elegans*, and geosmin has been proposed to serve as a non-toxic, chemical deterrent or aposematic signal (49). Carotenoids are lipophilic pigments produced by myxobacteria to quench toxic, reactive oxygen species (23, 25, 50). Iso-fatty acid signals produced by the VEPE/AEPE/TG-1 cluster are developmentally regulated signals that activate sporulation and fruiting body formation (27, 51, 52). Myxochelins are iron-chelating siderophores advantageous to iron competition in the environment (28, 29, 53-55). Notably, none of the metabolites with elucidated ecological roles are toxins such as antibacterials or antifungals; they instead have defensive, developmental, and metabolic roles. Although *in vitro* topoisomerase inhibition has been observed from alkylpyrones produced by *M. xanthus*, alkylpyrones and related alkylquinones and alkylresorcinols produced by type III PKSs have demonstrated a range of non-toxic activities such as alternative electron carriers in mycobacteria and membrane components that provide antibiotic resistance to *Streptomyces griseus* (30, 56, 57).

We suggest the vertical inheritance of BGCs encoding for metabolites that broadly benefit myxobacterial resilience during environmental stress explains conservation and maintenance of these clusters. We also suggest the absence of broadly conserved toxins, often considered to benefit predation, supports specialization of clades and potential competition for nutrients in the phylum. The recently elucidated role of ambruticin in myxobacteria-myxobacteria competition provides an example of how specialization might benefit myxobacterial genera and species (58). The limited number of clusters conserved in the phylum also corroborates the reported correlation between taxonomic distance and metabolic profiles of myxobacteria (9). Conservation of the horizontally-acquired indigoidine BGC in *Melittangium* offers insight into how gene transfer events could benefit specialization. Comparing the indigoidine clusters from *Str. chromofuscus* and *Str. aureofaciens*, the BpsA NRPS module includes a terminal tautomerase domain in the *Str. chromofuscus* cluster that is not present in BpsA from *Str. aureofaciens* (59). Instead, a discrete gene encoding for a homologous tautomerase is present in the indigoidine cluster from *Str. chromofuscus*. This observation suggests evolution of two, distinct indigoidine clusters that involved either excision of an NRPS-fused tautomerase or a fusion event between neighboring tautomerase and NPRS features. Mirroring these differences, the indigoidine BGC present in *Po*. sp. y55×33 includes a BpsA NRPS with a fused tautomerase, and the clusters from *Cystobacter* and *Melittangium* spp. have discrete tautomerase-encoding genes. Although parallel evolution of tautomerase fused/unfused indigoidine clusters in *Streptomyces* and myxobacteria cannot be ruled out, separate horizontal transfer events likely account for the differences in myxobacterial indigoidine BGCs. Our previous observation that the myxovirescin BGC conserved in *Myxococcus* spp. was acquired horizontally provides another example of a horizontal gene transfer event benefiting myxobacterial predation and specialization (16). Ultimately, our data and identification of conserved BGCs in Myxococcota reveals the vertical inheritance of biosynthetic pathways that produce metabolites involved in stress responses and chemical signaling, and the absence of BGCs encoding for toxic metabolites conserved at the phylum level supports competitive specialization of myxobacterial genera.

## Methods

### Isolation of myxobacteria

Environmental isolates were obtained using our previously established methodology (16, 17). Standard prey-baiting methods using *Escherichia coli* were used to isolate bacteriolytic myxobacteria. Air-dried soil samples were wetted with an antifungal solution (250 μg/mL of cycloheximide and nystatin) and pea-sized aliquots were plated on *E. coli* WAT plates. To prepare *E. coli* WAT plates an *E. coli* lawn was grown overnight at 37°C and resuspended in 1 mL of antifungal solution. Subsequently, 300 μl of the solution was spread across a WAT agar (1.5% agar, 0.1% CaCl_2_) plate and air-dried to yield *E. coli* WAT plates. Soil-inoculated plates were incubated at 25°C for up to 4 weeks with daily checks for the appearance of lytic zones or fruiting bodies, and observed lytic zones or fruiting bodies were passaged to VY/4 plates (Baker’s yeast 2.5 g/L, CaCl_2_ × 2H_2_O 1.36 g/L, vitamin B_12_ 0.5 mg/L, and agar 15 g/L) repeatedly until monocultures were obtained. Filter paper methodology was used to isolate cellulolytic myxobacteria as previously described. Briefly, individual squares of autoclaved filter paper were placed on ST21 agar plates (1 g/L of K_2_HPO_4_, 20 mg/L of yeast extract, 14 g/L of agar, 1 g/L of KNO_3_, 1 g/L of MgSO_4 ×_ 7H_2_O, 1 g/L of CaCl_2_ × 2H_2_O, 0.1 g/L of MnSO_4 ×_ 7H_2_O, and 0.2 g/L of FeCl_3_), and aliquots of antifungal solution wetted soil were placed on filter paper edges. Soil-inoculated plates were incubated at 25°C for up to 2 months with bidaily checks for growth starting after 2 weeks. Observed fruiting bodies were passaged to fresh ST21 filter paper plates until monocultures were obtained.

### Cultivation of isolates

All isolates were maintained on VY/4 plates and liquid cultures with CYH/2 media (0.75 g/L of casitone, 0.75g/L of yeast extract, 2g/L of starch, 0.5 g/L of soy flour, 0.5 g/L of glucose, 0.5 g/L of MgSO_4_•7H_2_O, 1 g/L of CaCl_2_•2H_2_O, 6 g/L of HEPES, 8 mg/L of EDTA-Fe, and 0.5 mg/L of vitamin B_12_).

### Sequencing methods

Monocultures of environmental isolates were utilized for isolation of genomic DNA using NucleoBond High Molecular Weight DNA Kits (Macherey-Nagel), and quality and concentration were assessed via Nanodrop spectrophotometry (Thermo Scientific NanoDrop One) and Qubit fluorometry (dsDNA HS Assay Kit, ThermoFisher Scientific). All isolates sequenced using an Oxford Nanopore Minion flow cell (R10.4.1) and associated ligation sequencing kit, native barcoding kit, or rapid barcoding kit. Basecalling and demultiplexing was performed using Guppy (v6+), assembly was performed using Flye (v2.9.2+), assembly error correction was performed using medaka (v1.11+) (60).

### Pan-genome analysis

Sequenced myxobacterial isolates and FASTA files for all *Anaeromyxobacter, Archangium, Corallococcus, Cystobacter, Melittangium, Myxococcus, Nannocystis, Polyangium, Pyxidicoccus, Sorangium*, and *Stigmatella* genomes available at NCBI Genome database were utilized for pan-genome analyses. All included myxobacteria can be found in Supplemental Table S1. Following established methodology, all genomes were annotated with Prokka (v1.11) to provide GFF3 files for pan-genome analysis using Roary (v3.12.0) (61-63). Pan-genomes were obtained for each analyzed genera and resulting gene_presence_absence.csv files were utilized for BGC analysis.

### BGC analysis

AntiSMASH version 7.0 was used to annotate BGCs present in all analyzed myxobacterial genomes (64). Genes included in resulting BGC Genbank files were subsequently used for comparison with gene presence/absence results from our pan-genome analysis to determine conserved genes within BGCs. For our analysis we considered conserved BGCs from each genus to be clusters present in 50% of the analyzed genomes (per genus) that included ≥2 conserved genes. All conserved genes observed in each of the 5 conserved BGCs can be found in Supplemental Tables S2-S7. Antismash analysis was also used for validation of identified, conserved BGCs from genera not included in our pan-genome analysis. Generated Genbank files from AntiSMASH were utilized as input in clinker on the CAGECAT webserver (v1.0) to produce BGC comparisons in Figures 2-5 (31, 33).

### Comparative genomics

OrthoANI calculations and tree generation were achieved using OAT (orthoANI tool v0.93.1) (32), and dDDH calculations were performed on the type strain genome server (TYGS) website (65).

### Heterologous production of indigoidine

The presence of an *indC*/*bpsA* homolog (WP_204491470.1) within the chromosome of *Me. primigenium* ATCC 29037 was observed by AntiSMASH analysis. Genomic DNA was extracted from *Me. primigenium* as previously described in the Sequencing methods section. Gene-specific primers were designed to amplify *indC*/*bpsA* from *Me. primigenium* and *Str. lavendulae* including 20–40 bp overlaps homologous to the multiple cloning sites of pET-28a to enable seamless Gibson Assembly and Circular Polymerase Extension Cloning (CPEC) based cloning (66). Primer sequences used for amplification and cloning listed in (Supplemental Table S8) were designed using SnapGene 7.1.1 and synthesized by Integrated DNA Technologies. PCR amplification of *indC*/*bpsA* from both *Me. primigenium* and, *Str. lavendulae*, was carried out using the TaKaRa PCR Amplification Kit. PCR products were purified using the GeneJET PCR Purification Kit (Thermo Fisher Scientific), and concentrations were determined via Nanodrop. Vector and insert DNA concentrations were normalized and combined at recommended molar ratios (2:1 insert to vector) according to the manufacturer’s protocol. Assembly of *indC*/*bpsA* from *Me. primigenium* into pET-28a was performed using Gibson Assembly (NEBuilder® HiFi DNA Assembly Master Mix), generating construct pNS001. The control construct, pNS002, containing *indC*/*bpsA* from *Str. lavendulae*, was assembled via CPEC using Platinum™ SuperFi II PCR Master Mix. Assembled plasmids were introduced into either DH10B or DH5α *E. coli* strains. Transformants were selected on LB agar with kanamycin (50 µg/mL). Plasmids were isolated from overnight cultures using the ZymoPURE II Plasmid Prep Kit. Linearization was performed by restriction digestion for length verification, followed by analysis via gel electrophoresis. Sequence confirmation of pNS001 and pNS002 was performed by Plasmidsaurus. Verified pNS001 and pNS002 constructs (Supplemental Table S9) were then transformed into *E. coli* BAP1 cells via electroporation for expression.

A single colony from each construct was inoculated in LB broth with kanamycin (50 μg/ml) and grown at 30°C to OD_600_ 0.4 before induction with 200 mM IPTG. Cultures were incubated overnight at 18 °C with shaking at 200 rpm. Indigoidine production was indicated by the development of blue pigmentation in liquid cultures. Following induction and cultivation, cultures were stored at −20 °C until subsequent LC-MS/MS analysis. Aliquots (1.5 mL) of overnight *E. coli* cultures exhibiting blue pigmentation were vortexed and centrifuged at 15,000 × g for 5 minutes to pellet the cells along with insoluble pigment. The pellet was resuspended in 1:1 methanol:water, vortexed briefly, and centrifuged again under the same conditions. The supernatant was discarded, and the pellet was resuspended in dimethyl sulfoxide (DMSO) to extract indigoidine. A 100 µL aliquot of this extract was then diluted with 900 µL of DMSO, resulting in a 1:10 dilution. The diluted extract was transferred to autosampler vials for LC-MS/MS analysis. Chromatographic and mass spectrometric analyses were performed using an Agilent 6530C Quadrupole Time-of-Flight (Q-TOF) mass spectrometer equipped with a Dual Agilent Jet Stream (AJS) electrospray ionization (ESI) source, interfaced with an Agilent 1260 Infinity II HPLC system. Chromatographic separation was carried out using a Gemini 5 μm NX-C18 110 Å, 50 × 2.0 mm analytical column (Phenomenex) at a constant flow rate of 0.5 mL/min. Mobile phase A consisted of water with 0.1% formic acid, while mobile phase B consisted of methanol with 0.1% formic acid. The gradient program began with 90% A and 10% B, followed by a linear increase to 25% A and 75% B at minute 5. The composition was then increased linearly to 10% A and 90% B over 1 minute and maintained at that mixture for an additional minute. Subsequently, the solvent composition was returned to the starting ratio over 1 minute. To ensure system stability and readiness for subsequent injections, a 2-minute re-equilibration phase was included. The total run time was set to 10 minutes, and the pressure limit was capped at 275 bar. A sample volume of 10 μL was injected per run. Injection draw and ejection speeds were set to 200 μL/min, and a needle height offset of 2 mm was used. A flush port needle wash (50% IPA/water) was enabled for 3 seconds, repeated three times. The autosampler was configured for sample overlap reduction with a sample flush-out factor of 5x the injection volume, though overlap injection mode was disabled for these runs. Data storage thresholds were set at 200 for MS and 5 for MS/MS, with centroid data collection enabled. The total cycle time was 2.3 seconds. The Q-TOF was operated in positive ionization mode with the following ion source conditions: gas temperature at 325 °C, drying gas flow at 10 L/min, nebulizer pressure at 50 psi, sheath gas temperature at 350 °C, and sheath gas flow at 12 L/min. Capillary voltage was set to 4000 V, with nozzle voltage at 0 V. Fragmentor voltage was 200 V, skimmer at 65 V, and octopole RF set to 750 V. Data were collected in centroid mode with MS scans acquired across a mass range of 100–1600 *m/z* at a rate of 5 spectra/s, and MS/MS scans across 50–1305 *m/z* at a rate of 3 spectra/s. Transient times were set to 200 ms/spectrum for MS and 333.3 ms/spectrum for MS/MS, with 2650 and 4355 transients/spectrum, respectively. Collision energies for MS/MS were calculated using the equation: (3 × *m/z* / 100) + 15. The system automatically selected precursor ions with a maximum of six precursors per cycle, applying active exclusion after three MS/MS spectra were collected per ion within a 0.2-minute window. Static exclusion windows were applied to exclude low- and high-mass ions (100–250 and 1300–1600 *m/z*) as well as known background ions, including *m/z* values of 922.0098, 531.40777, 553.38972, and 1083.791, with a delta tolerance of ±100 ppm. Precursor selection was based on abundance thresholds (10,000 counts, 0.01%) with iterative MS/MS enabled. Charge state preferences were set to prioritize 1+, 2+, and unknown precursors for fragmentation. Comparison of extracted ion chromatograms (EICs) for the exact mass of indigoidine (249-249.1 *m/z*) confirmed indigoidine production from *E. coli* BAP1 expressing *bpsA/indC* homologs from *Str. lavendulae* and *Me. primigenium*.

## Data availability statement

The datasets presented in this study can be found in online repositories. All NCBI accession number(s) can be found below:

ASM4919486v1 (BB12-2); ASM4919212v1 (hickory4); ASM4906018v1 (MIELM); ASM4906006v1 (NCWS); ASM4906002v1 (PVMSAZ); ASM4905998v1 (NCHinoki1); CP185339 (UBH4); CP185340 (D1P2); ASM4286526v1 (SCPoplar1)

## Supplementary Materials

Supplementary materials include: myxobacteria and genomes utilized for pan-genome analysis (Table S1); conserved features from carotenoid BGCs (Table S2), geosmin BGCs (Table S3), VEPE/AEPE/TG-1 BGCs (Table S4), myxochelin BGCs (Table S5), alkylpyrone BGCs (Table S6), and unknown type I PKS BGCs (Table S7); and primers used for *bpsA* cloning (Table S8)

## Funding

This research was supported by funds from the National Institute of General Medical Sciences (R01GM149795 and P20GM130460).

## Author Contributions

Isolation, cultivation, and sequencing of environmental myxobacteria A.A. and T.K.; pan-genome and BGC analysis S.K.; heterologous production of indigoidine N.S.; mass spectrometry N.S. and P.D.B.; conceptualization, manuscript preparation, and editing S.K., N.S., C.B.B., P.D.B., and D.C.S.; supervision and administration D.C.S. All authors have read and approved the final manuscript.

## Acknowledgements

The authors would like to thank Benjamin Pharr and the Mississippi Center for Supercomputing Research for their assistance with pan-genome analysis. The authors also utilized and appreciate the Glycoscience Center of Research Excellence Imaging Research Core.

## Notes

### Competing Interest Statement

The authors have declared no competing interest.

