## Supplemental for "Establishing Conserved Biosynthetic Gene Clusters of the Phylum Myxococcota"

**Supplemental Information**

**Table S1: Genomes included in pan-genome analysis.**

| ***Anaeromyxobacter*** | **Accession #** |
|---|---|
| *An. dehalogens 2CP1* | GCA_000022145.1 |
| *An. dehalogens 2CP-C* | GCA_000013385.1 |
| *An. sp. Fw109-5* | GCA_000017505.1 |
| *An. sp. K* | GCA_000020805.1 |
| *An. oryzae Red232* | GCA_023169945.1 |
| *An. paludicola Red630* | GCA_023169965.1 |
| *An. soli SG29* | GCA_022808855.1 |
| *An. diazotrophicus Red267* | GCA_013340205.1 |
| *An. sp. PSR-1* | GCA_000964525.1 |
| *An. sp. SG17* | GCA_022690695.1 |
| *An. terrae SG22* | GCA_022690685.1 |
| *An. sp. SG26* | GCA_022690725.1 |
| *An. soli SG29* | GCA_022808855.1 |
| *An. oryzisoli SG63* | GCA_022690765.1 |
| ***Archangium*** | |
| *Ar. lipolyticum* | GCA_024623785.1 |
| *Ar. sp. Cb G35* | GCA_001896145.1 |
| *Ar. violaceum Cb vi76* | GCA_003387095.1 |
| *Ar. gephyra DSM2261* | GCA_003387095.1 |
| *Ar. gephyra DSM2261v2* | GCA_001027285.1 |
| *Ar. lansingense* | GCA_026626635.1 |
| *Ar. lansignense NCHinoki1* | GCA_049059985.1 |
| *Ar. gephyra PVMSAZ* | GCA_049060025.1 |
| *Ar. violaceum scpoplar1* | GCA_042865265.1 |
| *Ar. violaceum SDU8* | GCA_016887565.1 |
| *Ar. violaceum SDU34* | GCA_016859125.1 |
| ***Corallococcus*** | |
| *Co. exiguus DSM14696* | GCA_006376655.1 |
| *Co. sp. AB011P* | GCA_003611605.1 |
| *Co. exiguus AB016* | GCA_012985275.1 |
| *Co. sp. AB018* | GCA_003986945.1 |
| *Co. sp. AB030* | GCA_003668965.1 |

| | |
|---|---|
| *Co. exiguus AB031* | GCA_012985225.1 |
| *Co. exiguus AB032A* | GCA_013155175.1 |
| *Co sp. AB032C* | GCA_003668935.1 |
| *Co. exiguus AB038A* | GCA_013155195.1 |
| *Co. sp. AB038B* | GCA_003668955.1 |
| *Co. exiguus AB039A* | GCA_013155575.1 |
| *Co. excercitus AB043A* | GCA_003611585.1 |
| *Co. excercitus AB043B* | GCA_013116705.1 |
| *Co. sp. AB045* | GCA_003668895.1 |
| *Co. interemptor AB047A* | GCA_003668875.1 |
| *Co. sp. AB049A* | GCA_003668885.1 |
| *Co. aberystwythensis AB050A* | GCA_003612165.1 |
| *Co. exiguus AM006* | GCA_013248865.1 |
| *Co. exiguus AM007* | GCA_013248925.1 |
| *Co. sp. AS-1-12* | GCA_020036995.1 |
| *Co. sp. AS-1-6* | GCA_020037015.1 |
| *Co. macrosporus ATCC29039* | GCA_017302985.1 |
| *Co. coralloides B035* | GCA_004104415.1 |
| *Co. sp. BB11-1* | GCA_026626625.1 |
| *Co. silvisoli C25j21* | GCA_009909145.1 |
| *Co. praedator CA031B* | GCA_003612125.1 |
| *Co. sp. CA031C* | GCA_003612115.1 |
| *Co. sicarius CA040B* | GCA_003611735.1 |
| *Co. sp. CA041A* | GCA_003612075.1 |
| *Co. carmarthensis CA043D* | GCA_003611695.1 |
| *Co. exercitus CA046A* | GCA_013116615.1 |
| *Co. carmarthensis CA046B* | GCA_013116625.1 |
| *Co. exiguus CA046D* | GCA_013248915.1 |
| *Co. sp. CA047B* | GCA_003612065.1 |
| *Co. exiguus CA048* | GCA_012985235.1 |
| *Co. sp. CA049B* | GCA_003611685.1 |
| *Co. llansteffanensis CA051B* | GCA_003612055.1 |
| *Co. sp. CA053C* | GCA_003611675.1 |
| *Co. terminator CA054A* | GCA_003611635.1 |
| *Co. sp. CA054B* | GCA_003611625.1 |
| *Co. exiguus DSM14696* | GCA_009909105.1 |
| *Co. macrosporus DSM14697* | GCA_002305895.1 |
| *Co. coralloides DSM2259* | GCA_000255295.1 |
| *Co. sp. EGB* | GCA_019968905.1 |

| | |
|---|---|
| *Co. macrosporus HW-1* | GCA_000219105.1 |
| *Co. exiguus NCCRE002* | GCA_017302975.1 |
| *Co. coralloides NCRR* | GCA_026965535.1 |
| *Co. sp. NCSPR001* | GCA_017309135.1 |
| *Co. sp. Z5C101001* | GCA_007352635.1 |
| *Co. soli ZKHCc11396* | GCA_014930455.1 |
| *Co. sp. bb12-1* | GCA_026626765.1 |
| ***Cystobacter*** | |
| *Cy. ferrugineus Cbfe23* | GCA_001887355.1 |
| *Cy. gracilis DSM14753* | GCA_020103725.1 |
| *Cy. fuscus DSM2262* | GCA_000335475.2 |
| *Cy. fuscus DSM52655* | GCA_002305875.1 |
| *Cy. fuscus NCWS* | GCA_049060065.1 |
| ***Melittangium*** | |
| *ATCC29037* | GCA_016904885.1 |
| *D1P2* | CP185340 |
| *DSM14713* | GCA_002305855.1 |
| *TKBC04* | - |
| ***Myxococcus*** | |
| *My. sp. AB022* | GCA_006547345.1 |
| *My. xanthus AB023* | GCA_013116805.1 |
| *My. sp. AB025B* | GCA_006518215.1 |
| *My. sp. AB036A* | GCA_006547355.1 |
| *My. eversor AB053B* | GCA_010894455.1 |
| *My. sp. AB056* | GCA_006547365.1 |
| *My. xanthus AM003* | GCA_013116825.1 |
| *My. xanthus AM005* | GCA_013116865.1 |
| *My. sp. AM009* | GCA_013372605.1 |
| *My. sp. AM010* | GCA_013372585.1 |
| *My. sp. AM011* | GCA_013372595.1 |
| *My. vastator AM301* | GCA_010894475.1 |
| *My. llanfairpwllgwyngyllgogerychwyrndrobwllllantysiliogogogochensis AM401* | GCA_006636215.1 |
| *My. sp. AS-1-15* | GCA_020037055.1 |
| *My. xanthus ATCC27925* | GCA_019895115.1 |
| *My. sp. BB12* | ASM4919486v1 |
| *My. sp. CA005* | GCA_006518205.1 |
| *My. sp. CA006* | GCA_006518195.1 |
| *My. sp. CA010* | GCA_006547325.1 |

| | |
|---|---|
| *My. sp. CA018* | GCA_010998655.1 |
| *My. sp. CA023* | GCA_010998615.1 |
| *My. sp. CA027* | GCA_010279825.1 |
| *My. xanthus CA029* | GCA_013116835.1 |
| *My. sp. CA033* | GCA_013336625.1 |
| *My. sp. CA039A* | GCA_013336645.1 |
| *My. sp. CA040A* | GCA_013336725.1 |
| *My. sp. CA051A* | GCA_013336705.1 |
| *My. sp. CA056* | GCA_013336715.1 |
| *My. stipitatus CYD_1* | GCA_021412625.1 |
| *My. xanthus DK101* | GCA_025739225.1 |
| *My. xanthus DK1050* | GCA_025739275.1 |
| *My. xanthus DK1622* | GCA_000012685.1 |
| *DK1622_Tpase* | GCA_015775755.1 |
| *My. stipitatus DSM14675* | GCA_000331735.1 |
| *My. fulvus DSM16525* | GCA_900111765.1 |
| *My. xanthus DSM16526* | GCA_900106535.1 |
| *My. virescens DSM2260* | GCA_900101905.1 |
| *My. xanthus DZ2(1)* | GCA_000278585.2 |
| *My. xanthus DZ2(2)* | GCA_018517205.1 |
| *My. xanthus DZ2(3)* | GCA_020827275.1 |
| *My. xanthus DZF1* | GCA_000340515.1 |
| *My. xanthus R31* | GCA_016698685.1 |
| *My. xanthus MC359c15* | GCA_006402735.1 |
| *My. xanthus MC335c16* | GCA_006402415.1 |
| *My. xanthus KF4.3.9c1* | GCA_006402015.1 |
| *My. xanthus GH3_5_6c2* | GCA_006400955.1 |
| *My. xanthus GH5_1_9c20* | GCA_006401215.1 |
| *My. fulvus Hickory4* | GCA_049192125.1 |
| *My. dinghuensis K15C18031901* | GCA_024198235.1 |
| *My. fulvus NBRC100333* | GCA_007991095.1 |
| *My. virescens NBRC100334* | GCA_007989405.1 |
| *My. qinghaiensis QH3KD-4-1* | GCA_024198215.1 |
| *My. fulvus 11* | GCA_023195975.1 |
| *My. guangdongensis K38C18041901* | GCA_024198255.1 |
| *My. xanthus KF3_28c_11* | GCA_006401635.1 |
| *My. hansupus* | GCA_000280925.3 |
| *My. sp. MISCRS1* | GCA_026626605.1 |
| *My. xanthus MxC21-1* | GCA_032612255.1 |

| | |
|---|---|
| *My. sp. NMCA1* | GCA_026810205.1 |
| *My. sp. RHSTA-1-4* | GCA_020037125.1 |
| *My. landrumensis SCHIC003* | GCA_017301635.1 |
| *My. sp. SDU36* | GCA_030168875.1 |
| *My. sp. XM-1-1* | GCA_020037095.1 |
| ***Nannocystis*** | |
| *N. exedens ATCC25963* | GCA_900112715.1 |
| *N. bainbridgea BB15-2* | GCA_028368995.1 |
| *Na. pusilla DSM53165* | GCA_020073745.1 |
| *Na. exedens DSM71* | GCA_002343915.1 |
| *Na. punicea FL3* | GCA_026965555.1 |
| *Na. sp. ILAH1* | GCA_026626585.1 |
| *Na. pusilla MIELM* | GCA_049060185.1 |
| *Na. radixulma NCELM* | GCA_028369095.1 |
| *Na. pusilla Na p29* | GCA_026626665.1 |
| *Na. sp. RBIL2* | GCA_026626745.1 |
| *Na. sp. SCPEA4* | GCA_026626685.1 |
| *Na. sp. UBH4* | CP185339 |
| ***Polyangium*** | |
| *Po. sp. 15x6* | GCA_029960785.1 |
| *Po. sp. 6x1* | GCA_029946515.1 |
| *Po. fumosum DSM14668* | GCA_005144585.1 |
| *Po. sorediatum DSM14670* | GCA_029946465.1 |
| *Po. spumosum DSM14734* | GCA_009649845.1 |
| *Po. mundeleinium RJM3* | GCA_028369105.1 |
| *Po. jinanense SDU13* | GCA_028435265.1 |
| *Po. jinanense SDU14* | GCA_028435365.1 |
| *Po. aurulentum SDU3-1* | GCA_005144635.2 |
| *Po. sp. y55x31* | GCA_029946505.1 |
| ***Pyxidicoccus*** | |
| *Py. caerfyrddinensis CA032A* | GCA_010894405.1 |
| *Py. fallax CA059B* | GCA_013155555.1 |
| *Py. trucidator CA060A* | GCA_010894435.1 |
| *Py. fallax DSM14698* | GCA_012933655.1 |
| *Py. sp. MSG2* | GCA_026626705.1 |
| *Py. xibeiensis QH1ED-7-1* | GCA_024198175.1 |
| *Py. parkwayensis SCPEA02* | GCA_017301735.1 |
| ***Sorangium*** | |
| *So. cellulosum So0007-03* | GCA_001589215.1 |

| | |
|---|---|
| *So. cellulosum So0008-312* | GCA_001589285.1 |
| *So. cellulosum So0011-07* | GCA_001589185.1 |
| *So. cellulosum So0149* | GCA_001589205.1 |
| *So. cellulosum So0157-18* | GCA_001589195.1 |
| *So. cellulosum So0157_2* | GCA_000418325.1 |
| *So. cellulosum So0157_25* | GCA_001589265.1 |
| *So. sp. Soce836* | GCA_028553905.1 |
| *So. cellulosum Soce26* | GCA_002950945.1 |
| *So. cellulosum Soce56* | GCA_000067165.1 |
| *So. cellulosum Soce836* | GCA_004135755.1 |
| *So. cellulosum SoceGT47* | GCA_004135735.1 |
| *So. atrum wiwo2* | GCA_028368935.1 |
| ***Stigmatella*** | |
| *St. hybrida DSM14722* | GCA_020103775.1 |
| *St. erecta DSM16858* | GCA_900111745.1 |
| *St. aurantiaca DSM17044* | GCA_900109545.1 |
| *St. aurantica DW4_3-1(p)* | GCA_000165485.1 |
| *St. aurantiaca DW4_3-1(c)* | GCA_000168055.1 |
| *St. ashevillensis ncwal01* | GCA_028368975.1 |

**Table S2. Carotenoid BGC conserved features**

| carotenoid | conserved gene | # of strains | # in BGC |
|---|---|---|---|
| **Archangium** | Dehydrosqualene desaturase | 12 | 12 |
| | hypothetical protein | 12 | 12 |
| | All-trans-phytoene synthase | 12 | 12 |
| | Polyketide biosynthesis 3-hydroxy-3-methylglutaryl-ACP synthase PksG | 12 | 12 |
| | zeta-carotene-forming phytoene desaturase | 12 | 12 |
| | Serine/threonine-protein kinase PknL | 12 | 12 |
| | Sensor protein FixL | 12 | 12 |
| | Putative acetolactate synthase large subunit IlvB2 | 10 | 10 |
| | S-(hydroxymethyl)glutathione dehydrogenase | 12 | 9 |
| | HTH-type transcriptional repressor YcgE | 9 | 9 |
| | Oxygen-independent coproporphyrinogen-III oxidase 1 | 9 | 9 |
| | HTH-type transcriptional repressor YcgE | 9 | 9 |
| | Oxygen-independent coproporphyrinogen-III oxidase 1 | 9 | 9 |
| | Serine/threonine-protein kinase Pkn1 | 9 | 9 |

| | | | |
|---|---|---|---|
| | hypothetical protein | 9 | 9 |
| | dTDP-4-amino-4,6-dideoxy-D-glucose transaminase | 9 | 9 |
| | hypothetical protein | 9 | 9 |
| | hypothetical protein | 9 | 9 |
| | S-formylglutathione hydrolase YeiG | 9 | 9 |
| | HTH-type transcriptional regulator AcrR | 9 | 9 |
| | Putative fatty-acid--CoA ligase fadD21 | 9 | 9 |
| | hypothetical protein | 9 | 9 |
| | hypothetical protein | 8 | 7 |
| | HTH-type transcriptional repressor YcgE | 7 | 7 |
| | hypothetical protein | 8 | 8 |
| | Acyclic carotenoid 1,2-hydratase | 7 | 7 |
| | hypothetical protein | 7 | 6 |
| | putative metallophosphoesterase | 7 | 7 |
| | NADH-quinone oxidoreductase subunit M | 6 | 6 |
| | Epimerase family protein | 6 | 6 |
| | hypothetical protein | 9 | 9 |
| | (2E,6E)-farnesyl diphosphate synthase | 6 | 6 |
| | Serine/threonine-protein kinase PknB | 6 | 6 |
| | HTH-type transcriptional regulator DmlR | 6 | 6 |
| | hypothetical protein | 6 | 6 |
| | hypothetical protein | 6 | 6 |
| | Na(+)/H(+) antiporter NhaG | 6 | 6 |
| | Methionine aminopeptidase 1, mitochondrial | 6 | 6 |
| | hypothetical protein | 6 | 6 |
| | hypothetical protein | 6 | 6 |
| | hypothetical protein | 6 | 6 |
| **Corallococcus** | zeta-carotene-forming phytoene desaturase | 49 | 48 |
| | All-trans-phytoene synthase | 39 | 39 |
| | Dehydrosqualene desaturase | 39 | 39 |
| | Acyclic carotenoid 1,2-hydratase | 37 | 37 |
| | hypothetical protein | 37 | 37 |

| | | | |
|---|---|---|---|
| | hypothetical protein | 35 | 34 |
| | hypothetical protein | 49 | 47 |
| | hypothetical protein | 49 | 39 |
| | Mercuric resistance operon regulatory protein | 33 | 31 |
| | HTH-type transcriptional repressor YcgE | 34 | 26 |
| | hypothetical protein | 29 | 25 |
| **Cystobacter** | Adenine deaminase | 5 | 5 |
| | hypothetical protein | 5 | 5 |
| | hypothetical protein | 5 | 5 |
| | Epimerase family protein | 5 | 5 |
| | hypothetical protein | 5 | 5 |
| | HTH-type transcriptional repressor YcgE | 5 | 5 |
| | hypothetical protein | 5 | 5 |
| | hypothetical protein | 5 | 5 |
| | Acyclic carotenoid 1,2-hydratase | 5 | 5 |
| | hypothetical protein | 5 | 5 |
| | Dehydrosqualene desaturase | 5 | 5 |
| | All-trans-phytoene synthase | 5 | 5 |
| | zeta-carotene-forming phytoene desaturase | 5 | 5 |
| | hypothetical protein | 5 | 4 |
| | putative metallophosphoesterase | 5 | 4 |
| | hypothetical protein | 5 | 4 |
| | Serine/threonine-protein kinase Pkn1 | 5 | 4 |
| | hypothetical protein | 3 | 3 |
| **Melittangium** | zeta-carotene-forming phytoene desaturase | 3 | 3 |
| | All-trans-phytoene synthase | 3 | 3 |
| | NADPH-dependent 7-cyano-7-deazaguanine reductase | 3 | 3 |
| | Serine/threonine-protein kinase PrkC | 3 | 3 |
| | Guanosine-5'-triphosphate,3'-diphosphate pyrophosphatase | 3 | 3 |
| | hypothetical protein | 3 | 3 |

| | | | |
|---|---|---|---|
| **Myxococcus** | HTH-type transcriptional repressor YcgE | 37 | 36 |
| | HTH-type transcriptional repressor YcgE | 38 | 37 |
| | hypothetical protein | 39 | 38 |
| | hypothetical protein | 39 | 38 |
| | hypothetical protein | 41 | 40 |
| | hypothetical protein | 38 | 37 |
| | Acyclic carotenoid 1,2-hydratase | 38 | 37 |
| | Dehydrosqualene desaturase | 39 | 38 |
| | All-trans-phytoene synthase | 39 | 38 |
| | zeta-carotene-forming phytoene desaturase | 39 | 38 |
| | hypothetical protein | 39 | 35 |
| | Epimerase family protein | 38 | 34 |
| | Enterochelin esterase | 39 | 35 |
| | hypothetical protein | 38 | 34 |
| **Polyangium** | Spore protein SP21 | 9 | 9 |
| | HTH-type transcriptional repressor YcgE | 9 | 9 |
| | Phytoene desaturase (lycopene-forming) | 9 | 9 |
| | Hydroxyneurosporene desaturase | 9 | 9 |
| | Farnesyl diphosphate synthase | 9 | 9 |
| | Alkaline phosphatase synthesis sensor protein PhoR | 9 | 6 |
| | hypothetical protein | 8 | 6 |
| | hypothetical protein | 9 | 7 |
| | Spore protein SP21 | 8 | 8 |
| | hypothetical protein | 8 | 8 |
| | hypothetical protein | 8 | 8 |
| | hypothetical protein | 8 | 8 |
| | Acyclic carotenoid 1,2-hydratase | 8 | 8 |
| | RsbT co-antagonist protein RsbRD | 9 | 9 |
| | 15-cis-phytoene synthase | 8 | 8 |
| **Pyxidicoccus** | Dehydrosqualene desaturase | 6 | 5 |
| | 3-hydroxy-3-methylglutaryl-coenzyme A reductase | 6 | 5 |

| | | | |
|---|---|---|---|
| | hypothetical protein | 7 | 6 |
| | zeta-carotene-forming phytoene desaturase | 7 | 7 |
| | hypothetical protein | 5 | 5 |
| | Epimerase family protein | 4 | 4 |
| | Acyclic carotenoid 1,2-hydratase | 4 | 4 |
| | hypothetical protein | 4 | 4 |
| | HTH-type transcriptional repressor YcgE | 4 | 4 |
| | hypothetical protein | 3 | 3 |
| | All-trans-phytoene synthase | 3 | 3 |
| **Sorangium** | Phytoene desaturase (lycopene-forming) | 14 | 8 |
| | hypothetical protein | 13 | 8 |
| | Farnesyl diphosphate synthase | 13 | 7 |
| **Stigmatella** | Octaprenyl-diphosphate synthase | 6 | 6 |
| | zeta-carotene-forming phytoene desaturase | 6 | 6 |
| | Acyclic carotenoid 1,2-hydratase | 6 | 6 |
| | hypothetical protein | 6 | 5 |
| | Dehydrosqualene desaturase | 3 | 3 |
| | All-trans-phytoene synthase | 3 | 3 |
| | hypothetical protein | 3 | 3 |
| | Sensor protein ZraS | 3 | 3 |
| | putative HTH-type transcriptional regulator YybR | 3 | 3 |
| | putative metallophosphoesterase | 3 | 3 |
| | hypothetical protein | 3 | 3 |
| | Mercuric resistance operon regulatory protein | 3 | 3 |
| | Sensor protein ZraS | 3 | 3 |
| | Epimerase family protein | 3 | 3 |
| | hypothetical protein | 3 | 3 |
| | All-trans-phytoene synthase | 3 | 3 |
| | 3-hydroxy-3-methylglutaryl-coenzyme A reductase | 3 | 3 |
| | Dehydrosqualene desaturase | 3 | 3 |
| | hypothetical protein | 3 | 3 |

| | | | |
|---|---|---|---|
| | hypothetical protein | 3 | 3 |
| | HTH-type transcriptional repressor YcgE | 3 | 3 |
| | Extracellular serine proteinase precursor | 3 | 3 |

**Table S3. Geosmin BGC conserved features**

| geosmin | conserved gene | # of strains | # in BGC |
|---|---|---|---|
| **Archangium** | Major membrane protein I | 9 | 9 |
| | Germacradienol/geosmin synthase | 7 | 7 |
| | Major membrane protein I | 7 | 7 |
| | hypothetical protein | 9 | 6 |
| | Protease HtpX | 9 | 6 |
| | N-acetylneuraminate epimerase | 6 | 6 |
| **Corallococcus** | Rod shape-determining protein MreB | 50 | 25 |
| | Undecaprenyl-phosphate mannosyltransferase | 51 | 27 |
| | L-2,4-diaminobutyrate decarboxylase | 46 | 27 |
| | 2,3,4,5-tetrahydropyridine-2,6-dicarboxylate N-succinyltransferase | 51 | 40 |
| | Succinyl-diaminopimelate desuccinylase | 49 | 40 |
| | hypothetical protein | 37 | 30 |
| | Germacradienol/geosmin synthase | 39 | 32 |
| | Major membrane protein I | 49 | 49 |
| | Major membrane protein I | 39 | 39 |
| | hypothetical protein | 25 | 25 |
| | Cytochrome c-type protein NrfH | 28 | 27 |
| | Cytochrome c-552 precursor | 31 | 28 |
| | hypothetical protein | 36 | 33 |
| **Cystobacter** | 4'-demethylrebeccamycin synthase | 5 | 5 |
| | Germacradienol/geosmin synthase | 5 | 5 |
| | hypothetical protein | 5 | 5 |
| | Major membrane protein I | 5 | 5 |
| | Major membrane protein I | 5 | 5 |
| | UDP-glucose 6-dehydrogenase TuaD | 5 | 5 |

| | | | |
|---|---|---|---|
| | Serine/threonine-protein kinase StkP | 5 | 3 |
| | Multidrug resistance protein 3 | 4 | 4 |
| | hypothetical protein | 3 | 3 |
| **Melittangium** | zeta-carotene-forming phytoene desaturase | 3 | 3 |
| | All-trans-phytoene synthase | 3 | 3 |
| | NADPH-dependent 7-cyano-7-deazaguanine reductase | 3 | 3 |
| | Serine/threonine-protein kinase PrkC | 3 | 3 |
| | Guanosine-5'-triphosphate,3'-diphosphate pyrophosphatase | 3 | 3 |
| | hypothetical protein | 3 | 3 |
| **Myxococcus** | Germacradienol/geosmin synthase | 36 | 35 |
| | Transposon Tn10 TetC protein | 39 | 34 |
| | hypothetical protein | 39 | 35 |
| | hypothetical protein | 38 | 34 |
| **Nannocystis** | Major membrane protein I | 12 | 11 |
| | Germacradienol/geosmin synthase | 11 | 11 |
| | Major membrane protein I | 9 | 8 |
| | putative MscS family protein YkuT | 8 | 7 |
| | Minor extracellular protease Epr precursor | 7 | 6 |
| **Polyangium** | Major membrane protein I | 9 | 9 |
| | Germacradienol/geosmin synthase | 8 | 8 |
| | L-glyceraldehyde 3-phosphate reductase | 8 | 8 |
| | HTH-type transcriptional repressor ComR | 8 | 8 |
| | Major membrane protein I | 8 | 8 |
| **Pyxidicoccus** | Germacradienol/geosmin synthase | 5 | 5 |
| | Major membrane protein I | 5 | 5 |
| | Major membrane protein I | 5 | 5 |
| | putative oxidoreductase | 6 | 3 |

| | | | |
|---|---|---|---|
| **Sorangium** | Germacradienol/geosmin synthase | 8 | 8 |
| | Major membrane protein I | 8 | 8 |
| | Major membrane protein I | 8 | 8 |
| | putative MscS family protein YkuT | 7 | 7 |
| **Stigmatella** | Spermidine synthase | 6 | 4 |
| | hypothetical protein | 6 | 4 |
| | hypothetical protein | 6 | 4 |
| | NADP-dependent alcohol dehydrogenase C 2 | 3 | 3 |
| | Major membrane protein I | 3 | 3 |
| | hypothetical protein | 3 | 3 |
| | F420-dependent glucose-6-phosphate dehydrogenase | 3 | 3 |
| | Serine/threonine-protein kinase pkn6 | 3 | 3 |
| | hypothetical protein | 3 | 3 |
| | hypothetical protein | 3 | 3 |
| | HTH-type transcriptional regulator DmlR | 3 | 3 |
| | Major membrane protein I | 3 | 3 |
| | Germacradienol/geosmin synthase | 3 | 3 |
| | Major membrane protein I | 3 | 3 |
| | Germacradienol/geosmin synthase | 3 | 3 |
| | Phosphatidylglycerol lysyltransferase | 3 | 3 |
| | Sporulation initiation phosphotransferase F | 3 | 3 |
| | Major membrane protein I | 3 | 3 |

**Table S4. VEPE/AEPE/TG-1 BGC conserved features**

| **VEPE/AEPE/TG-1** | **conserved gene** | **# of strains** | **# in BGC** |
|---|---|---|---|
| **Anaeromyxobacter** | | | |
| **Archangium** | Threonylcarbamoyl-AMP synthase | 12 | 6 |
| | Protein ApaG | 12 | 12 |
| | Glycogen debranching enzyme | 12 | 12 |
| | hypothetical protein | 12 | 12 |
| | Phosphoserine phosphatase | 12 | 12 |
| | hypothetical protein | 12 | 12 |

| | | | |
|---|---|---|---|
| | Long-chain-fatty-acid--CoA ligase | 12 | 12 |
| | 3 beta-hydroxysteroid dehydrogenase/Delta 5-->4-isomerase | 12 | 12 |
| | Response regulator PleD | 12 | 10 |
| | Glucose-1-phosphate adenylyltransferase | 12 | 11 |
| | Aminodeoxyfutalosine deaminase | 12 | 11 |
| | RNA-splicing ligase RtcB | 11 | 9 |
| | Ferrochelatase | 10 | 10 |
| | 1D-myo-inositol 2-acetamido-2-deoxy-alpha-D-glucopyranoside deacetylase | 7 | 7 |
| | Maltooligosyl trehalose synthase | 7 | 7 |
| | Inner membrane protein alx | 9 | 9 |
| | Putative ketoacyl reductase | 7 | 7 |
| | hypothetical protein | 6 | 6 |
| | 3 beta-hydroxysteroid dehydrogenase/Delta 5-->4-isomerase | 6 | 6 |
| **Corallococcus** | Outer membrane protein assembly factor BamB precursor | 49 | 34 |
| | Aminodeoxyfutalosine deaminase | 51 | 41 |
| | Glucose-1-phosphate adenylyltransferase | 51 | 41 |
| | hypothetical protein | 51 | 42 |
| | Phosphoserine phosphatase | 49 | 36 |
| | Glycogen debranching enzyme | 49 | 25 |
| | Protein ApaG | 49 | 25 |
| | Putative niacin/nicotinamide transporter NaiP | 39 | 29 |
| | hypothetical protein | 36 | 27 |
| | 3 beta-hydroxysteroid dehydrogenase/Delta 5-->4-isomerase | 49 | 38 |
| | Long-chain-fatty-acid--CoA ligase | 40 | 40 |
| | 3 beta-hydroxysteroid dehydrogenase/Delta 5-->4-isomerase | 49 | 28 |
| | Glycogen debranching enzyme | 49 | 25 |
| **Cystobacter** | Response regulator rcp1 | 5 | 5 |
| | Glucose-1-phosphate adenylyltransferase | 5 | 5 |
| | hypothetical protein | 5 | 5 |
| | hypothetical protein | 5 | 5 |

| | | | |
|---|---|---|---|
| | Phosphoserine phosphatase | 5 | 5 |
| | Ferrochelatase | 5 | 5 |
| | Bacteriophytochrome | 5 | 5 |
| | hypothetical protein | 5 | 5 |
| | ATP-dependent DNA helicase PcrA | 5 | 5 |
| | 3 beta-hydroxysteroid dehydrogenase/Delta 5-->4-isomerase | 5 | 5 |
| | Glycogen debranching enzyme | 5 | 5 |
| | Inner membrane protein alx | 5 | 5 |
| | hypothetical protein | 5 | 5 |
| | hypothetical protein | 5 | 5 |
| | hypothetical protein | 5 | 5 |
| | hypothetical protein | 5 | 5 |
| | Multidrug resistance protein NorM | 5 | 5 |
| | Aminodeoxyfutalosine deaminase | 5 | 5 |
| | Glucose 1-dehydrogenase 4 | 5 | 5 |
| | Response regulator PleD | 5 | 5 |
| | Maltooligosyl trehalose synthase | 5 | 5 |
| | Protein ApaG | 5 | 5 |
| | Threonylcarbamoyl-AMP synthase | 5 | 5 |
| | Epimerase family protein | 5 | 5 |
| | hypothetical protein | 5 | 5 |
| | 3 beta-hydroxysteroid dehydrogenase/Delta 5-->4-isomerase | 5 | 5 |
| | Long-chain-fatty-acid--CoA ligase | 5 | 5 |
| | Bis(5'-nucleosyl)-tetraphosphatase PrpE [asymmetrical] | 4 | 4 |
| **Melittangium** | Phosphoserine phosphatase | 4 | 3 |
| | Response regulator PleD | 4 | 3 |
| | Glycogen debranching enzyme | 4 | 3 |
| | Threonylcarbamoyl-AMP synthase | 4 | 3 |
| | Protein ApaG | 4 | 3 |
| | Inner membrane protein alx | 4 | 3 |
| | hypothetical protein | 4 | 3 |
| | Long-chain-fatty-acid--CoA ligase | 4 | 3 |

| | | | |
|---|---|---|---|
| | 3 beta-hydroxysteroid dehydrogenase/Delta 5-->4-isomerase | 4 | 3 |
| | Glucose-1-phosphate adenylyltransferase | 4 | 3 |
| | Aminodeoxyfutalosine deaminase | 4 | 3 |
| | ATP-dependent DNA helicase PcrA | 4 | 3 |
| | hypothetical protein | 4 | 3 |
| | Response regulator rcp1 | 4 | 3 |
| **Myxococcus** | Ferrochelatase | 39 | 37 |
| | Protein ApaG | 62 | 59 |
| | Inner membrane protein alx | 39 | 37 |
| | Maltooligosyl trehalose synthase | 39 | 37 |
| | Glycogen debranching enzyme | 39 | 37 |
| | Phosphoserine phosphatase | 62 | 61 |
| | Long-chain-fatty-acid--CoA ligase | 39 | 37 |
| | hypothetical protein | 63 | 61 |
| | Levodione reductase | 39 | 37 |
| | Response regulator PleD | 62 | 59 |
| | 3 beta-hydroxysteroid dehydrogenase/Delta 5-->4-isomerase | 39 | 36 |
| | GDP-6-deoxy-D-mannose reductase | 38 | 36 |
| | hypothetical protein | 38 | 36 |
| | RNA-splicing ligase RtcB | 38 | 37 |
| | hypothetical protein | 38 | 35 |
| | Outer membrane protein assembly factor BamB precursor | 38 | 35 |
| | hypothetical protein | 62 | 50 |
| | hypothetical protein | 38 | 36 |
| | putative N-acetyltransferase YafP | 37 | 34 |
| | hypothetical protein | 62 | 52 |
| | Aminodeoxyfutalosine deaminase | 63 | 60 |
| | hypothetical protein | 63 | 61 |
| | Glucose-1-phosphate adenylyltransferase | 63 | 60 |
| **Pyxidicoccus** | 3 beta-hydroxysteroid dehydrogenase/Delta 5-->4-isomerase | 7 | 7 |
| | GDP-6-deoxy-D-mannose reductase | 7 | 7 |

| | | | |
|---|---|---|---|
| | Long-chain-fatty-acid--CoA ligase | 7 | 7 |
| | hypothetical protein | 7 | 7 |
| | Phosphoserine phosphatase | 7 | 7 |
| | Glycogen debranching enzyme | 7 | 6 |
| | hypothetical protein | 7 | 6 |
| | Protein ApaG | 7 | 5 |
| | Glucose-1-phosphate adenylyltransferase | 7 | 5 |
| | RNA-splicing ligase RtcB | 7 | 5 |
| | hypothetical protein | 7 | 5 |
| | Response regulator PleD | 7 | 5 |
| | Aminodeoxyfutalosine deaminase | 7 | 5 |
| | hypothetical protein | 5 | 5 |
| | 4-formylbenzenesulfonate dehydrogenase TsaC1/TsaC2 | 5 | 5 |
| | ATP-dependent DNA helicase PcrA | 5 | 5 |
| | hypothetical protein | 7 | 4 |
| | Maltooligosyl trehalose synthase | 4 | 4 |
| | hypothetical protein | 4 | 4 |
| | putative peptidase | 3 | 3 |
| | Ferrochelatase | 3 | 3 |
| | Inner membrane protein alx | 3 | 3 |
| | hypothetical protein | 3 | 3 |
| | Threonylcarbamoyl-AMP synthase | 7 | 3 |
| **Stigmatella** | 3-oxoacyl-[acyl-carrier-protein] reductase FabG | 6 | 6 |
| | Glycogen debranching enzyme | 6 | 6 |
| | Inner membrane protein alx | 6 | 6 |
| | Ferrochelatase | 6 | 6 |
| | Threonylcarbamoyl-AMP synthase | 6 | 6 |
| | hypothetical protein | 6 | 6 |
| | hypothetical protein | 6 | 6 |
| | hypothetical protein | 6 | 6 |
| | ATP-dependent DNA helicase PcrA | 6 | 6 |
| | Glucose-1-phosphate adenylyltransferase | 6 | 6 |

| | | | |
|---|---|---|---|
| | 3 beta-hydroxysteroid dehydrogenase/Delta 5-->4-isomerase | 6 | 6 |
| | Long-chain-fatty-acid--CoA ligase | 6 | 6 |
| | Protein ApaG | 6 | 6 |
| | Aminodeoxyfutalosine deaminase | 6 | 6 |
| | Response regulator PleD | 6 | 6 |
| | hypothetical protein | 6 | 6 |
| | Phosphoserine phosphatase | 6 | 6 |
| | putative ABC transporter ATP-binding protein YxlF | 6 | 6 |
| | hypothetical protein | 6 | 6 |
| | hypothetical protein | 6 | 6 |
| | putative sensor histidine kinase TcrY | 3 | 3 |
| | hypothetical protein | 3 | 3 |
| | Metallo-beta-lactamase L1 precursor | 3 | 3 |
| | hypothetical protein | 3 | 3 |
| | hypothetical protein | 3 | 3 |
| | hypothetical protein | 3 | 3 |
| | Membrane-bound lytic murein transglycosylase A precursor | 3 | 3 |
| | Maltooligosyl trehalose synthase | 3 | 3 |
| | Epimerase family protein | 3 | 3 |
| | hypothetical protein | 3 | 3 |
| | hypothetical protein | 6 | 3 |
| | Membrane-bound lytic murein transglycosylase A precursor | 3 | 3 |
| | Hydroxyacylglutathione hydrolase | 3 | 3 |
| | Maltooligosyl trehalose synthase | 3 | 3 |
| | Epimerase family protein | 3 | 3 |

#### Table S5. Myxochelin BGC conserved features

| myxochelin | conserved gene | # of strains | # in BGC |
|---|---|---|---|
| **Archangium** | 2,3-dihydro-2,3-dihydroxybenzoate dehydrogenase | 11 | 11 |
| | hypothetical protein | 12 | 9 |
| | hypothetical protein | 11 | 9 |
| | hypothetical protein | 9 | 9 |
| | hypothetical protein | 9 | 9 |

| | | | |
|---|---|---|---|
| | hypothetical protein | 10 | 8 |
| | hypothetical protein | 8 | 8 |
| | hypothetical protein | 8 | 8 |
| | Phospho-2-dehydro-3-deoxyheptonate aldolase | 8 | 8 |
| | HTH-type transcriptional regulator TtgR | 9 | 7 |
| | Putative ribosome biogenesis GTPase RsgA | 7 | 6 |
| | 2,3-dihydroxybenzoate-AMP ligase | 6 | 6 |
| | Isochorismate synthase DhbC | 6 | 6 |
| | hypothetical protein | 6 | 6 |
| | Dimodular nonribosomal peptide synthase | 6 | 6 |
| | hypothetical protein | 6 | 6 |
| | Isochorismatase | 6 | 6 |
| | Pentalenene oxygenase | 6 | 6 |
| **Corallococcus** | hypothetical protein | 39 | 26 |
| | Phospho-2-dehydro-3-deoxyheptonate aldolase | 46 | 45 |
| | Dimodular nonribosomal peptide synthase | 36 | 35 |
| | Isochorismatase | 35 | 32 |
| | 2,3-dihydroxybenzoate-AMP ligase | 44 | 42 |
| | Isochorismate synthase DhbC | 36 | 32 |
| | 2,3-dihydro-2,3-dihydroxybenzoate dehydrogenase | 36 | 31 |
| | 3-aminobutyryl-CoA aminotransferase | 48 | 40 |
| | Purine efflux pump PbuE | 36 | 30 |
| **Cystobacter** | Biopolymer transport protein ExbB | 5 | 5 |
| | cAMP receptor protein | 5 | 5 |
| | HTH-type transcriptional repressor KstR2 | 5 | 5 |
| | 3-oxoadipate CoA-transferase subunit A | 5 | 5 |
| | Dimodular nonribosomal peptide synthase | 5 | 5 |
| | 2,3-dihydroxybenzoate-AMP ligase | 5 | 5 |
| | Serine acetyltransferase | 5 | 5 |
| | hypothetical protein | 5 | 5 |
| | Phospho-2-dehydro-3-deoxyheptonate aldolase | 5 | 5 |

| | | | |
|---|---|---|---|
| | Isochorismatase | 5 | 5 |
| | Isochorismate synthase DhbC | 5 | 5 |
| | 2,3-dihydro-2,3-dihydroxybenzoate dehydrogenase | 5 | 5 |
| | Thiol-disulfide oxidoreductase ResA | 5 | 5 |
| | hypothetical protein | 5 | 5 |
| | Thioredoxin reductase | 5 | 5 |
| | O-succinylhomoserine sulfhydrylase | 5 | 5 |
| | Cyclic pyranopterin monophosphate synthase accessory protein 2 | 5 | 5 |
| | Cysteine synthase | 5 | 5 |
| | Biopolymer transport protein ExbD | 5 | 5 |
| | Alkaline phosphatase synthesis transcriptional regulatory protein PhoP | 5 | 5 |
| | 3-oxoadipate CoA-transferase subunit B | 5 | 4 |
| | Alcohol dehydrogenase | 5 | 4 |
| | hypothetical protein | 4 | 4 |
| | hypothetical protein | 4 | 4 |
| | hypothetical protein | 4 | 4 |
| | hypothetical protein | 4 | 4 |
| | hypothetical protein | 4 | 4 |
| | Vitamin B12 transporter BtuB precursor | 4 | 4 |
| | 6-phosphogluconolactonase | 5 | 3 |
| | hypothetical protein | 5 | 3 |
| | Polyketide biosynthesis 3-hydroxy-3-methylglutaryl-ACP synthase PksG | 4 | 3 |
| | hypothetical protein | 4 | 3 |
| | Sensor protein kinase WalK | 4 | 4 |
| | hypothetical protein | 3 | 3 |
| | hypothetical protein | 3 | 3 |
| **Melittangium** | 50S ribosomal protein L13 | 4 | 4 |
| | 30S ribosomal protein S9 | 4 | 4 |
| | FHA domain-containing protein FhaB | 4 | 4 |
| | Selenide, water dikinase | 4 | 4 |
| | Ribonuclease PH | 4 | 4 |

| | | | |
|---|---|---|---|
| | hypothetical protein | 4 | 3 |
| | Twitching mobility protein | 4 | 3 |
| | ATP-dependent DNA helicase PcrA | 4 | 3 |
| | hypothetical protein | 4 | 3 |
| | hypothetical protein | 4 | 3 |
| | hypothetical protein | 4 | 3 |
| | hypothetical protein | 4 | 3 |
| | hypothetical protein | 4 | 3 |
| | Mannan endo-1,4-beta-mannosidase precursor | 4 | 3 |
| | NTE family protein RssA | 4 | 3 |
| | hypothetical protein | 3 | 3 |
| | hypothetical protein | 3 | 3 |
| | Regulatory protein RecX | 3 | 3 |
| | Outer membrane protein assembly factor BamD precursor | 3 | 3 |
| | N-acetylmuramoyl-L-alanine amidase AmiC precursor | 3 | 3 |
| | Non-canonical purine NTP pyrophosphatase | 3 | 3 |
| | hypothetical protein | 3 | 3 |
| | hypothetical protein | 3 | 3 |
| | Aminopeptidase S | 3 | 3 |
| | hypothetical protein | 3 | 3 |
| | hypothetical protein | 3 | 3 |
| | ABC transporter ATP-binding protein YojI | 3 | 3 |
| | Potassium-transporting ATPase A chain | 3 | 3 |
| | Potassium-transporting ATPase B chain | 3 | 3 |
| | Sensor protein KdpD | 3 | 3 |
| | hypothetical protein | 3 | 3 |
| | Biotin biosynthesis cytochrome P450 | 3 | 3 |
| | hypothetical protein | 3 | 3 |
| | Demethylrebeccamycin-D-glucose O-methyltransferase | 3 | 3 |
| | NADP-dependent alcohol dehydrogenase C 2 | 3 | 3 |
| | hypothetical protein | 3 | 3 |
| | Linear gramicidin synthase subunit D | 3 | 3 |
| | cAMP receptor protein | 3 | 3 |

| | | | |
|---|---|---|---|
| | hypothetical protein | 3 | 3 |
| | Tyrocidine synthase 3 | 3 | 3 |
| | Pentalenene oxygenase | 3 | 3 |
| | hypothetical protein | 3 | 3 |
| | Potassium-transporting ATPase C chain | 3 | 3 |
| | Alginate biosynthesis sensor protein KinB | 3 | 3 |
| | hypothetical protein | 3 | 3 |
| | Polyketide synthase PksL | 3 | 3 |
| | Polyketide biosynthesis protein PksE | 3 | 3 |
| | Pentachlorophenol 4-monooxygenase | 3 | 3 |
| | Polyketide synthase PksJ | 3 | 3 |
| | hypothetical protein | 3 | 3 |
| | hypothetical protein | 3 | 3 |
| | Serine/threonine-protein kinase Pkn1 | 3 | 3 |
| | Polyketide biosynthesis 3-hydroxy-3-methylglutaryl-ACP synthase PksG | 3 | 3 |
| | Alcohol dehydrogenase | 3 | 3 |
| | 3-oxoadipate CoA-transferase subunit B | 3 | 3 |
| | 3-oxoadipate CoA-transferase subunit A | 3 | 3 |
| | HTH-type transcriptional repressor KstR2 | 3 | 3 |
| | Cyclic pyranopterin monophosphate synthase accessory protein 2 | 3 | 3 |
| | DNA protection during starvation protein | 3 | 3 |
| | Isochorismate synthase DhbC | 3 | 3 |
| | Serine acetyltransferase | 3 | 3 |
| | Cysteine synthase | 3 | 3 |
| | Prolyl tripeptidyl peptidase precursor | 3 | 3 |
| | hypothetical protein | 3 | 3 |
| **Myxococcus** | hypothetical protein | 48 | 35 |
| | Macrolide export ATP-binding/permease protein MacB | 38 | 36 |
| | 2,3-dihydro-2,3-dihydroxybenzoate dehydrogenase | 38 | 37 |
| | Isochorismate synthase DhbC | 38 | 37 |
| | 2,3-dihydroxybenzoate-AMP ligase | 38 | 37 |

| | | | |
|---|---|---|---|
| | Isochorismatase | 38 | 37 |
| | Dimodular nonribosomal peptide synthase | 38 | 37 |
| | Phospho-2-dehydro-3-deoxyheptonate aldolase | 38 | 37 |
| | Hexuronate transporter | 38 | 37 |
| | 3-aminobutyryl-CoA aminotransferase | 38 | 37 |
| | Vibriobactin utilization protein ViuB | 38 | 37 |
| | hypothetical protein | 38 | 37 |
| | hypothetical protein | 38 | 33 |
| | Limonene 1,2-monooxygenase | 38 | 33 |
| | Phthiocerol/phenolphthiocerol synthesis polyketide synthase type I PpsE | 37 | 32 |
| | Phthiocerol/phenolphthiocerol synthesis polyketide synthase type I PpsE | 37 | 32 |
| | Glycogen synthase | 38 | 32 |
| | Linear gramicidin dehydrogenase LgrE | 38 | 32 |
| | hypothetical protein | 38 | 32 |
| | hypothetical protein | 38 | 32 |
| | hypothetical protein | 38 | 32 |
| | hypothetical protein | 37 | 32 |
| | hypothetical protein | 38 | 32 |
| | Phthiocerol synthesis polyketide synthase type I PpsE | 36 | 32 |
| **Polyangium** | Vitamin B12 transporter BtuB precursor | 7 | 7 |
| | hypothetical protein | 7 | 7 |
| | Glutamate-1-semialdehyde 2,1-aminomutase | 7 | 7 |
| | Phospho-2-dehydro-3-deoxyheptonate aldolase | 5 | 5 |
| **Pyxidicoccus** | cAMP receptor protein | 7 | 5 |
| | hypothetical protein | 7 | 4 |
| | Bifunctional protein PaaZ | 6 | 4 |
| | Isochorismate synthase DhbC | 4 | 4 |
| | 2,3-dihydro-2,3-dihydroxybenzoate dehydrogenase | 4 | 4 |
| | Vibriobactin utilization protein ViuB | 3 | 3 |
| | D-alanine--D-alanine ligase | 4 | 3 |

| | | | |
|---|---|---|---|
| | Phospho-2-dehydro-3-deoxyheptonate aldolase | 3 | 3 |
| | 2,3-dihydroxybenzoate-AMP ligase | 3 | 3 |
| | Adenylate cyclase 2 | 3 | 3 |
| | Putative outer membrane protein precursor | 3 | 3 |
| | Putative pyridoxal phosphate-dependent acyltransferase | 3 | 3 |
| | Phthiocerol synthesis polyketide synthase type I PpsC | 3 | 3 |
| | Heptaprenyl diphosphate synthase component 2 | 7 | 3 |
| | Biopolymer transport protein ExbB | 4 | 4 |
| | Quaternary ammonium compound-resistance protein SugE | 7 | 3 |
| | Phthiocerol synthesis polyketide synthase type I PpsE | 3 | 3 |
| **Stigmatella** | Isochorismatase | 6 | 6 |
| | Isochorismate synthase DhbC | 6 | 6 |
| | 2,3-dihydroxybenzoate-AMP ligase | 6 | 6 |
| | NADPH-dependent ferric-chelate reductase | 6 | 6 |
| | Transcription elongation factor GreB | 6 | 6 |
| | PhoH-like protein | 6 | 5 |
| | Biotin biosynthesis cytochrome P450 | 6 | 5 |
| | Hemin transport system permease protein HmuU | 6 | 5 |
| | 3-aminobutyryl-CoA aminotransferase | 6 | 5 |
| | Phospho-2-dehydro-3-deoxyheptonate aldolase | 6 | 5 |
| | hypothetical protein | 6 | 5 |
| | ATP-dependent RNA helicase HrpB | 4 | 3 |
| | hypothetical protein | 3 | 3 |
| | hypothetical protein | 3 | 3 |
| | Purine efflux pump PbuE | 3 | 3 |
| | Vitamin B12 transporter BtuB precursor | 3 | 3 |
| | 2,3-dihydro-2,3-dihydroxybenzoate dehydrogenase | 3 | 3 |
| | Pyridoxal 4-dehydrogenase | 3 | 3 |
| | hypothetical protein | 3 | 3 |
| | Trans-aconitate 2-methyltransferase | 3 | 3 |
| | hypothetical protein | 3 | 3 |
| | Hemin-binding periplasmic protein HmuT precursor | 3 | 3 |

| | | | |
|---|---|---|---|
| | Hemin import ATP-binding protein HmuV | 3 | 3 |
| | hypothetical protein | 3 | 3 |
| | Dimodular nonribosomal peptide synthase | 3 | 3 |
| | Phthiotriol/phenolphthiotriol dimycocerosates methyltransferase | 3 | 3 |
| | hypothetical protein | 3 | 3 |
| | Peptide methionine sulfoxide reductase MsrB | 3 | 3 |
| | hypothetical protein | 3 | 3 |
| | hypothetical protein | 6 | 4 |
| | lipid kinase YegS | 6 | 4 |
| | putative oxidoreductase | 6 | 4 |
| | Chaperone protein DnaK | 6 | 4 |
| | Dimodular nonribosomal peptide synthase | 3 | 3 |
| | 2,3-dihydro-2,3-dihydroxybenzoate dehydrogenase | 3 | 3 |
| | hypothetical protein | 3 | 3 |
| | 3-isopropylmalate dehydrogenase | 3 | 3 |
| | 3-isopropylmalate dehydratase small subunit | 3 | 3 |
| | 3-isopropylmalate dehydratase large subunit | 3 | 3 |
| | 2-isopropylmalate synthase | 3 | 3 |
| | hypothetical protein | 3 | 3 |
| | Ubiquinone/menaquinone biosynthesis C-methyltransferase UbiE | 6 | 3 |
| | hypothetical protein | 4 | 3 |

**Table S6. Alkylpyrone BGC conserved features**

| **alkylpyrone** | **conserved gene** | **# of strains** | **# in BGC** |
|---|---|---|---|
| **Archangium** | Putative peroxiredoxin bcp | 11 | 11 |
| | Response regulator SaeR | 12 | 11 |
| | 2-octaprenyl-3-methyl-6-methoxy-1,4-benzoquinol hydroxylase | 12 | 11 |
| | Alpha-pyrone synthesis polyketide synthase-like Pks11 | 12 | 11 |
| | hypothetical protein | 12 | 11 |
| | Multifunctional cyclase-dehydratase-3-O-methyl transferase TcmN | 12 | 11 |
| | hypothetical protein | 12 | 10 |
| | putative lipoprotein YbbD precursor | 12 | 10 |
| | hypothetical protein | 10 | 9 |

| | | | |
|---|---|---|---|
| | putative oxidoreductase | 12 | 9 |
| | hypothetical protein | 9 | 9 |
| | Acyl carrier protein | 9 | 9 |
| | hypothetical protein | 9 | 9 |
| | Putative ligase | 9 | 9 |
| | Alpha-pyrone synthesis polyketide synthase-like Pks18 | 9 | 9 |
| | Methyl-accepting chemotaxis protein CtpH | 9 | 9 |
| | Carbonic anhydrase 1 | 9 | 8 |
| | hypothetical protein | 9 | 8 |
| | Bicarbonate transporter BicA | 9 | 8 |
| | hypothetical protein | 8 | 7 |
| | Long-chain-fatty-acid--AMP ligase FadD26 | 7 | 7 |
| | Decaprenyl-phosphate phosphoribosyltransferase | 7 | 7 |
| | hypothetical protein | 7 | 7 |
| | Luminescence regulatory protein LuxO | 7 | 6 |
| | hypothetical protein | 6 | 6 |
| | Cocaine esterase | 6 | 6 |
| | Meromycolate extension acyl carrier protein | 6 | 6 |
| | hypothetical protein | 6 | 6 |
| | Alkaline phosphatase synthesis transcriptional regulatory protein PhoP | 6 | 6 |
| | hypothetical protein | 6 | 6 |
| **Corallococcus** | Alpha-pyrone synthesis polyketide synthase-like Pks11 | 49 | 30 |
| | hypothetical protein | 49 | 30 |
| | Long-chain-fatty-acid--AMP ligase FadD26 | 49 | 30 |
| | hypothetical protein | 46 | 30 |
| | hypothetical protein | 47 | 28 |
| | hypothetical protein | 41 | 27 |
| | hypothetical protein | 39 | 27 |
| | hypothetical protein | 39 | 26 |
| | Multifunctional cyclase-dehydratase-3-O-methyl transferase TcmN | 49 | 28 |
| | putative transcriptional regulatory protein TcrX | 49 | 27 |

| | | | |
|---|---|---|---|
| | hypothetical protein | 49 | 25 |
| **Cystobacter** | Proline--tRNA ligase | 5 | 4 |
| | Alpha-pyrone synthesis polyketide synthase-like Pks18 | 5 | 4 |
| | hypothetical protein | 5 | 4 |
| | Beta-glucanase precursor | 5 | 4 |
| | Putative trans-acting enoyl reductase | 5 | 4 |
| | Methyl-accepting chemotaxis protein CtpH | 5 | 4 |
| | Long-chain-fatty-acid--AMP ligase FadD29 | 5 | 4 |
| | Putative ligase/MSMEI_5285 | 5 | 4 |
| | High-affinity branched-chain amino acid transport ATP-binding protein LivF | 5 | 4 |
| | hypothetical protein | 5 | 4 |
| | hypothetical protein | 5 | 4 |
| | Endonuclease YhcR precursor | 5 | 4 |
| | hypothetical protein | 5 | 4 |
| | Alpha-pyrone synthesis polyketide synthase-like Pks11 | 5 | 4 |
| | Acyl carrier protein | 5 | 4 |
| | Acyl carrier protein | 5 | 4 |
| | putative decaprenylphosphoryl-beta-D-ribose oxidase | 5 | 4 |
| | Decaprenyl-phosphate phosphoribosyltransferase | 5 | 4 |
| | Multifunctional cyclase-dehydratase-3-O-methyl transferase TcmN | 5 | 4 |
| | putative transcriptional regulatory protein TcrX | 5 | 4 |
| | D-alanine--D-alanine ligase | 5 | 4 |
| | hypothetical protein | 5 | 4 |
| | hypothetical protein | 5 | 4 |
| | putative oxidoreductase | 5 | 4 |
| | hypothetical protein | 5 | 4 |
| **Melittangium** | Multifunctional cyclase-dehydratase-3-O-methyl transferase TcmN | 4 | 3 |
| | Acyl carrier protein | 4 | 3 |
| | Alpha-pyrone synthesis polyketide synthase-like Pks18 | 4 | 3 |
| | Long-chain-fatty-acid--AMP ligase FadD29 | 4 | 3 |

| | | | |
|---|---|---|---|
| | Alpha-pyrone synthesis polyketide synthase-like Pks11 | 4 | 3 |
| | Proline--tRNA ligase | 4 | 3 |
| | High-affinity branched-chain amino acid transport ATP-binding protein LivF | 4 | 3 |
| | Lipopolysaccharide export system ATP-binding protein LptB | 4 | 3 |
| | hypothetical protein | 4 | 3 |
| | Methyl-accepting chemotaxis protein CtpH | 4 | 3 |
| **Myxococcus** | Circadian clock protein kinase KaiC | 38 | 34 |
| | Sporulation initiation phosphotransferase F | 39 | 35 |
| | putative lipoprotein YbbD precursor | 39 | 35 |
| | Putative peroxiredoxin bcp | 39 | 35 |
| | hypothetical protein | 38 | 34 |
| | hypothetical protein | 39 | 36 |
| | putative transcriptional regulatory protein TcrX | 64 | 46 |
| | hypothetical protein | 38 | 35 |
| | Multifunctional cyclase-dehydratase-3-O-methyl transferase TcmN | 39 | 36 |
| | Decaprenyl-phosphate phosphoribosyltransferase | 39 | 36 |
| | putative decaprenylphosphoryl-beta-D-ribose oxidase | 39 | 36 |
| | putative oxidoreductase | 63 | 56 |
| | hypothetical protein | 39 | 35 |
| | 3-(3-hydroxy-phenyl)propionate/3-hydroxycinnamic acid hydroxylase | 38 | 35 |
| | Long-chain-fatty-acid--AMP ligase FadD29 | 38 | 35 |
| | Meromycolate extension acyl carrier protein | 39 | 38 |
| | hypothetical protein | 38 | 37 |
| | Alpha-pyrone synthesis polyketide synthase-like Pks11 | 39 | 38 |
| | Sulfate/thiosulfate import ATP-binding protein CysA | 38 | 36 |
| | Molybdenum transport system permease protein ModB | 40 | 38 |
| | Molybdate-binding periplasmic protein precursor | 38 | 36 |
| | Organic hydroperoxide resistance transcriptional regulator | 38 | 36 |
| | hypothetical protein | 39 | 37 |
| | hypothetical protein | 37 | 36 |
| | hypothetical protein | 39 | 37 |

| | | | |
|---|---|---|---|
| | hypothetical protein | 38 | 36 |
| | hypothetical protein | 38 | 35 |
| | hypothetical protein | 37 | 34 |
| | hypothetical protein | 39 | 37 |
| | Proline--tRNA ligase | 62 | 40 |
| | hypothetical protein | 38 | 36 |
| | hypothetical protein | 38 | 36 |
| | hypothetical protein | 36 | 34 |
| | hypothetical protein | 36 | 33 |
| | hypothetical protein | 37 | 36 |
| | hypothetical protein | 37 | 34 |
| | hypothetical protein | 36 | 34 |
| | hypothetical protein | 36 | 33 |
| **Pyxidicoccus** | hypothetical protein | 7 | 5 |
| | 3-(3-hydroxy-phenyl)propionate/3-hydroxycinnamic acid hydroxylase | 7 | 5 |
| | Molybdenum transport system permease protein ModB | 7 | 5 |
| | Long-chain-fatty-acid--AMP ligase FadD29 | 7 | 5 |
| | Glucose-1-phosphate adenylyltransferase | 7 | 5 |
| | hypothetical protein | 7 | 5 |
| | Sulfate/thiosulfate import ATP-binding protein CysA | 5 | 5 |
| | Alpha-pyrone synthesis polyketide synthase-like Pks11 | 5 | 5 |
| | Multidrug resistance operon repressor | 4 | 4 |
| | hypothetical protein | 3 | 3 |
| | Molybdate-binding periplasmic protein precursor | 3 | 3 |
| | Transcriptional activator NphR | 3 | 3 |
| | Acyl carrier protein | 3 | 3 |
| **Stigmatella** | hypothetical protein | 6 | 6 |
| | putative oxidoreductase | 6 | 6 |
| | Alpha-pyrone synthesis polyketide synthase-like Pks11 | 6 | 6 |
| | putative transcriptional regulatory protein TcrX | 6 | 6 |

| | | | |
|---|---|---|---|
| | Multifunctional cyclase-dehydratase-3-O-methyl transferase TcmN | 6 | 6 |
| | Proline--tRNA ligase | 6 | 6 |
| | Putative peroxiredoxin bcp | 6 | 6 |
| | Meromycolate extension acyl carrier protein | 6 | 6 |
| | hypothetical protein | 6 | 6 |
| | hypothetical protein | 3 | 3 |
| | Long-chain-fatty-acid--AMP ligase FadD26 | 3 | 3 |
| | Decaprenyl-phosphate phosphoribosyltransferase | 3 | 3 |
| | putative decaprenylphosphoryl-beta-D-ribose oxidase | 3 | 3 |
| | Virulence sensor protein BvgS precursor | 3 | 3 |
| | 3-(3-hydroxy-phenyl)propionate/3-hydroxycinnamic acid hydroxylase | 3 | 3 |
| | putative HTH-type transcriptional regulator YusO | 3 | 3 |
| | hypothetical protein | 3 | 3 |
| | hypothetical protein | 3 | 3 |
| | 2'-5'-RNA ligase | 3 | 3 |
| | hypothetical protein | 3 | 3 |
| | hypothetical protein | 3 | 3 |
| | hypothetical protein | 3 | 3 |
| | hypothetical protein | 3 | 3 |
| | hypothetical protein | 3 | 3 |
| | putative HTH-type transcriptional regulator YusO | 3 | 3 |
| | Long-chain-fatty-acid--AMP ligase FadD26 | 3 | 3 |
| | 3-(3-hydroxy-phenyl)propionate/3-hydroxycinnamic acid hydroxylase | 3 | 3 |
| | putative decaprenylphosphoryl-beta-D-ribose oxidase | 3 | 3 |
| | Decaprenyl-phosphate phosphoribosyltransferase | 3 | 3 |
| | hypothetical protein | 3 | 3 |
| | hypothetical protein | 3 | 3 |

**Table S7. Unknown type I PKS BGC conserved features**

| type I PKS | conserved gene | # of strains | # in BGC |
|---|---|---|---|

| | | | |
|---|---|---|---|
| **Archangium** | Sporulation initiation phosphotransferase F | 12 | 11 |
| | Toluene 1,2-dioxygenase system ferredoxin subunit | 12 | 11 |
| | putative FAD-linked oxidoreductase | 12 | 11 |
| | RNA polymerase sigma factor SigA | 12 | 11 |
| | Acyl-CoA dehydrogenase | 12 | 11 |
| | hypothetical protein | 11 | 10 |
| | hypothetical protein | 12 | 8 |
| | hypothetical protein | 10 | 8 |
| | High-affinity zinc uptake system membrane protein ZnuB | 10 | 9 |
| | Erythronolide synthase, modules 1 and 2 | 9 | 8 |
| | GTP cyclohydrolase 1 | 9 | 8 |
| | Manganese ABC transporter substrate-binding lipoprotein precursor | 9 | 8 |
| | High-affinity zinc uptake system ATP-binding protein ZnuC | 9 | 8 |
| | Formyl-coenzyme A transferase | 9 | 8 |
| | hypothetical protein | 8 | 7 |
| | Long-chain-fatty-acid--CoA ligase FadD15 | 8 | 7 |
| | Non-motile and phage-resistance protein | 8 | 7 |
| | hypothetical protein | 7 | 6 |
| | hypothetical protein | 6 | 6 |
| **Melittangium** | Erythronolide synthase, modules 1 and 2 | 3 | 3 |
| | Phthiocerol synthesis polyketide synthase type I PpsC | 3 | 3 |
| | Acyl-CoA dehydrogenase | 3 | 3 |
| | hypothetical protein | 3 | 3 |
| | RNA polymerase sigma factor SigA | 3 | 3 |
| | putative FAD-linked oxidoreductase | 3 | 3 |
| | Zinc import ATP-binding protein ZnuC | 3 | 3 |
| | Manganese transport system membrane protein MntB | 3 | 3 |

### Table S8. Primer table

| Primer Name | Sequence (5' to 3') | Product Size (bp) | Purpose |
|---|---|---|---|

| | | | |
|---|---|---|---|
| S_bpsA_F | ctcaaactagataccaggcatccgaaaggaagctgagttggctg | 3971 | Amplify *S. lavendulae bpsA* with overlap for pet-28a vector |
| S_bpsA_R | catcgtcgtttcctcgcatcgtggcatggtatatctccttcttaaagttaaac | 3971 | Amplify *S. lavendulae bpsA* with overlap for pet-28a vector |
| S_pet-28a_F | ctcaaactagataccaggcatccgaaaggaagctgagttggctg | 5234 | Amplify pet-28a vector with overlap for *S. lavendulae bpsA* |
| S_pet-28a_R | *catcgtcgtttcctcgcatcgtggcatggtatatctccttcttaaagttaaac* | 5234 | Amplify pet-28a vector with overlap for *S. lavendulae bpsA* |
| MbpsA_V1_F | gtttaactttaagaaggagatataccatgaatacggaaattctggcgaaagc | 3966 | Amplify *M. primigenium bpsA* with overlap for pet-28a vector |
| MbpsA_V1_R | cagccaactcagcttcctttcgggatgtcgtggattagcattggc | 3966 | Amplify *M. primigenium bpsA* with overlap for pet-28a vector |
| M_pet-28a_v1_F | gtttaactttaagaaggagatataccatgaatacggaaattctggcgaaagc | 5234 | Amplify pet-28a vector with overlap for *M. primigenium bpsA* |
| M_pet-28a_v1_R | gctttcgccagaatttccgtattcatggtatatctccttcttaaagttaaac | 5234 | Amplify pet-28a vector with overlap for *M. primigenium bpsA* |
| M_bpsA_v2_F | gccaatgctaatccacgacatcccgaaaggaagctgagttggctg | 4105 | Amplify *M. primigenium bpsA* with overlap for BAC vector |
| M_bpsA_v2_R | tggtatctagtttgagtctgcggatgtcgtggattagcattggc | 4105 | Amplify *M. primigenium bpsA* with overlap for BAC vector |
| S_*bpsA*_check_F | gaagagcaagctccaggtcaagg | 958 | Check for the presence of *bpsA* from *S. lavendulae* |
| S_*bpsA*_check_R | gttcatcatcaggtgctggagcttc | 958 | Check for the presence of *bpsA* from *S. lavendulae* |
| M_*bpsA*_check_F | gatcgagaaccacgactgggtc | 1240 | Check for the presence of *bpsA* from *M. primigenium* |
| M_*bpsA*_check_R | gaagaagtaggagggaccgctctc | 1240 | Check for the presence of *bpsA* from *M. primigenium* |

**Table S9. Plasmid table**

| Plasmid name | Gene of interest | Vector | Antibiotic resistance | promotor | source |
|---|---|---|---|---|---|
| pNS001 | *bpsA* sourced from *M. primigenium* | pet-28a | kanR | T7 | This study |
| pNS002 | *bpsA* sourced from *S. lavendulae* | pet-28a | kanR | T7 | This study |
